# Deep learning super-resolution of paediatric ultra-low-field MRI without paired high-field scans

**DOI:** 10.1101/2024.11.29.625898

**Authors:** Ula Briski, Niall J. Bourke, Hajer Karoui, Kirsten A. Donald, Layla E. Bradford, Simone R. Williams, Michal R. Zieff, Sadia Parkar, Sidra Kaleem, Salman Osmani, Sean C.L. Deoni, Steven C.R. Williams, Khula South Africa Study Team, Rosalyn J. Moran, Levente Baljer, František Váša

## Abstract

Brain magnetic resonance imaging (MRI) is essential for diagnosis and neurodevelopmental research, but the high cost and infrastructure demands of high-field MRI limit its use to high-income settings. Ultra-low-field MRI scanners offer a more affordable and energy-efficient alternative, but their reduced resolution and signal-to-noise ratio restrict research and clinical utility, prompting the need for super-resolution techniques. Current super-resolution methods rely on either three anisotropic ultra-low-field scans acquired at different orientations (axial, coronal, sagittal) to reconstruct a higher-resolution image using multi-resolution registration (MRR) or the training of deep learning models using paired ultra-low- and high-field scans. Since acquiring three high-quality ultra-low-field scans is not always feasible, and paired high-field data may not be available, this study explores the efficacy of using a deep learning model to generate scans of MRR quality from a single ultra-low-field input scan. Results demonstrated significant enhancement in the quality of output scans, including improved image quality metrics, stronger tissue volume correlations, and greater Dice overlap of tissue segmentations. Generating higher-resolution brain scans from single ultra-low-field scans, without paired high-field data, reduces scanning time and further widens MRI accessibility in low- and middle-income countries. This approach also facilitates site-specific model training, which an exploratory external validation suggests may be necessary to address potential domain shifts across scanning sites.

## Introduction

Magnetic resonance imaging (MRI) is widely used for medical diagnosis and research [1]. Unlike other neuroimaging techniques, such as computed tomography and positron emission tomography, MRI is non-invasive and does not use ionizing radiation, making it ideal for studying, for example, foetal and early infant neurodevelopment [2]. Despite its utility, MRI technology is not equally accessible to everyone. In low- and middle-income countries (LMICs), fewer than one MRI scanner is available per million individuals [3]. For example, there are only 84 MRI scanners available to 373 million people in the West African sub-region [4]. Conversely, high-income countries (HICs) have about one MRI unit for every 25,000 people [4]. Moreover, MRI scanners in LMICs are frequently outdated or predominantly located in urban areas [5], further reducing their accessibility. The main reasons for this are the high capital equipment costs and infrastructural requirements [6]. These include purchasing the system, installing it in a shielded suite, powering it, the need for cryogenic cooling, and ongoing maintenance costs, as well as the need for trained specialists [7] [8]. In addition to significantly limiting clinical use, this disparity also negatively affects research efforts. For example, studies on neurodevelopment and neural correlates of disorders often rely on small sample sizes and are predominantly conducted on participants from HICs, thus limiting their generalizability [2] [9].

One possible way to address this problem is the adoption of ultra-low-field (ULF) MRI scanners that use substantially lower magnetic field strength (e.g. <0.1 T) than conventional high-field (HF) scanners (1.5-3 T). ULF scanners are considerably cheaper and easier to maintain, and additionally require less power and no specific facilities [6]. Therefore, these systems could offer better paediatric clinical care in LIMCs and opportunities for carrying out larger studies with diverse populations. Deoni and colleagues [2] demonstrated that a 64mT ULF MRI scanner could reliably replicate brain volume and developmental estimates from a 3T HF scanner, with additional benefits including quieter scan acquisition and allowing parents to be present, resulting in reduced movement artefacts. Despite the many benefits of ULF systems and the potential for their global adoption, the primary drawback is that resulting scans have significantly lower signal-to-noise ratio (SNR) and resolution compared to HF systems. For example, scans acquired with the 64 mT Hyperfine Swoop have a spatial resolution of 1.5 × 1.5 × 5 mm^3^ compared to standard 1 × 1 × 1 mm^3^ in HF scanners. This yields anisotropic images where the resolution is high within one plane (axial, coronal, or sagittal) with reasonably small pixels (1.5 × 1.5 mm^2^), but with considerably thicker slices (5 mm). Each anisotropic image takes 3-6 minutes to acquire, depending on contrast and SNR, making it particularly useful for infants who cannot keep still for long periods.

One approach to improve the utility of lower-quality ultra-low-field scans is super-resolution (SR). SR algorithms aim to increase the resolution and SNR of an image and their use has been strongly driven by the development of new deep learning techniques in recent years [10], primarily through the use of convolutional neural networks (CNN). For example, the UNet was originally developed for the segmentation of biomedical images but is now also widely used for SR problems [11] [12]. Iglesias and colleagues [13] introduced a 3D UNet super-resolution method for reconstruction of high-quality MRI brain scans from inputs of varied contrasts and resolutions, allowing for the analysis of a wide range of MRI data. This model, however, was trained on synthetically generated scans of healthy adults, hindering its applicability to paediatric data. Several other approaches have been proposed, including a multi-orientation UNet [14], generative adversarial networks (GANs) [15], and SRDenseNet [16]; however, they all require paired ULF and HF data for training. Beyond classical CNN-based architectures, attention-based models such as Attention UNet [17] and transformer-based architectures such as SwinUNETR [18] have recently emerged for medical image analysis tasks including segmentation and reconstruction. These architectures offer improved contextual modelling and may further benefit SR applications.

A study by Deoni and colleagues [19] presents a successful SR approach for reconstructing higher-resolution isotropic images from three low-resolution anisotropic images (axial, coronal, and sagittal) acquired at low magnetic field strengths, without the use of deep learning, called multi-resolution registration (MRR). MRR involves combining three ULF scans with dimensions of 1.5 × 1.5 × 5 mm^3^, each with high-resolution within one orthogonal plane (axial, coronal, sagittal), into a single 1.5 × 1.5 × 1.5 mm^3^ higher-resolution image. The MRR approach to SR, however, has the limitation of requiring acquisition of all three anisotropic ULF scans to reconstruct a higher-resolution isotropic scan. This results in a long scanning protocol (9-18 min) compared to the modern HF scanners. Moreover, it is not always possible to acquire good-quality scans from all three orientations due to head motion or other artefacts, particularly in paediatric populations.

To overcome this problem, this study investigates the effectiveness of deep learning models at reconstructing higher-resolution scans of MRR quality (1.5 × 1.5 × 1.5 mm^3^) from any anisotropic ULF scan (1.5 × 1.5 × 5 mm^3^) using T2-weighted 64 mT ULF scans of 6-month-old infants. Higher-resolution images were reconstructed using deep learning models without the need for paired ULF and HF data. The quality of super-resolved model outputs shows increased correlations of tissue volume and greater Dice overlap of segmented brain regions to MRR reference scans. Obtaining a 1.5 × 1.5 × 1.5 mm^3^ scan from a single anisotropic low-resolution scan can help reduce scanning time, making this approach especially useful for scanning neonates, infants and any other participants who struggle to remain still during lengthy scanner sessions. Additionally, less time in the scanner means lower costs, further widening accessibility to MRI in LMICs.

## Methods

### MRI Data

All procedures were approved by the Human Research Ethics Committee at the University of Cape Town in South Africa. Trained study staff obtained consent from the participants’ parents prior to participation in any study-specific procedures. The consent forms were translated into local languages, and participants were informed that participation is voluntary and that they are free to withdraw at any point without any repercussions. Families were compensated after each study visit. All experiments were performed in accordance with relevant named guidelines and regulations.

The MRI data used in the study was acquired in Cape Town, South Africa as part of the ‘Khula Study’ [20], which is a multi-site longitudinal birth cohort study investigating executive function emergence and development in the first 2 years of life in South Africa and Malawi. To be eligible for participation, women had to be in their third trimester of pregnancy or up to three months postpartum. Additionally, women had to be older than 18 years, had to have had a singleton pregnancy, and not use psychotropic drugs during pregnancy. Women with major complications during delivery or infants with the presence of congenital malformations or abnormalities were also excluded. In South Africa, researchers recruited 394 participants. A range of data types were collected, including demographic and health-related information, neuroimaging, electrophysiological and behavioural data, as well as biospecimens. MRI data was acquired at both ultra-low-field (ULF), using a 64 mT Hyperfine Swoop system, and at high-field (HF), using a 3T Siemens system. In the present study, T2-weighted ULF scans from 6-month-old infants were used. We only used data from a single age group as the contrast between grey and white matter changes significantly during the first two years of life due to rapid myelination of white matter [21] [22], and including multiple age groups could make the learning task more difficult.

Each ULF scan has high resolution within one plane – axial (AXI), coronal (COR) or sagittal (SAG) – but low resolution along the remaining dimension (i.e., 1.5 × 1.5 × 5 mm^3^). In the present study, we only included participants with all three orthogonal anisotropic scans (referred to as AXI, COR and SAG), yielding a sample size of 200 participants. We performed visual quality control (QC) to exclude any participants with at least one low-quality anisotropic ULF scan. Participants were removed from the dataset if any one of their three scans exhibited severe motion artefacts or if part of the brain was out of the field of view. As a result, 35 participants were excluded, leaving a final sample of 165 participants. See Supplementary Fig. S1 for examples of excluded scans.

### Pre-processing

We performed rigid registration of each participant’s ULF scans to a HF template using the SPM12 software package [23]. The template, serving as the reference image, was developed by Baljer and colleagues [14] from HF scans of age-corresponding participants. Following this step, an additional 33 participants were excluded due to failed registration, resulting in a final dataset of 132 participants, each contributing three ULF scans (a total of 396 scans). Following registration, the three anisotropic ULF scans (AXI, COR, SAG) from each participant were combined into a single higher-resolution MRR scan, using multi-resolution registration [19] (Fig. 1A). Similarly to Deoni and colleagues [19], we used the *antsMultivariateTemplateConstruction2.sh* function from the Advanced Normalization Tools (ANTs) software package [24] to repeatedly co-register and average the three ULF scans, finally generating a higher-resolution scan with an effective voxel resolution of 1.5 × 1.5 × 1.5 mm^3^. The resulting dataset comprised of three anisotropic scans (AXI, COR, SAG) and one corresponding MRR scan per participant that would later be used to sample paired data for training and evaluation of the SR model. Data were split into training/validation/test sets of 86/20/26 subjects (65/15/20 %) (Fig. 1A).

**Figure 1:**
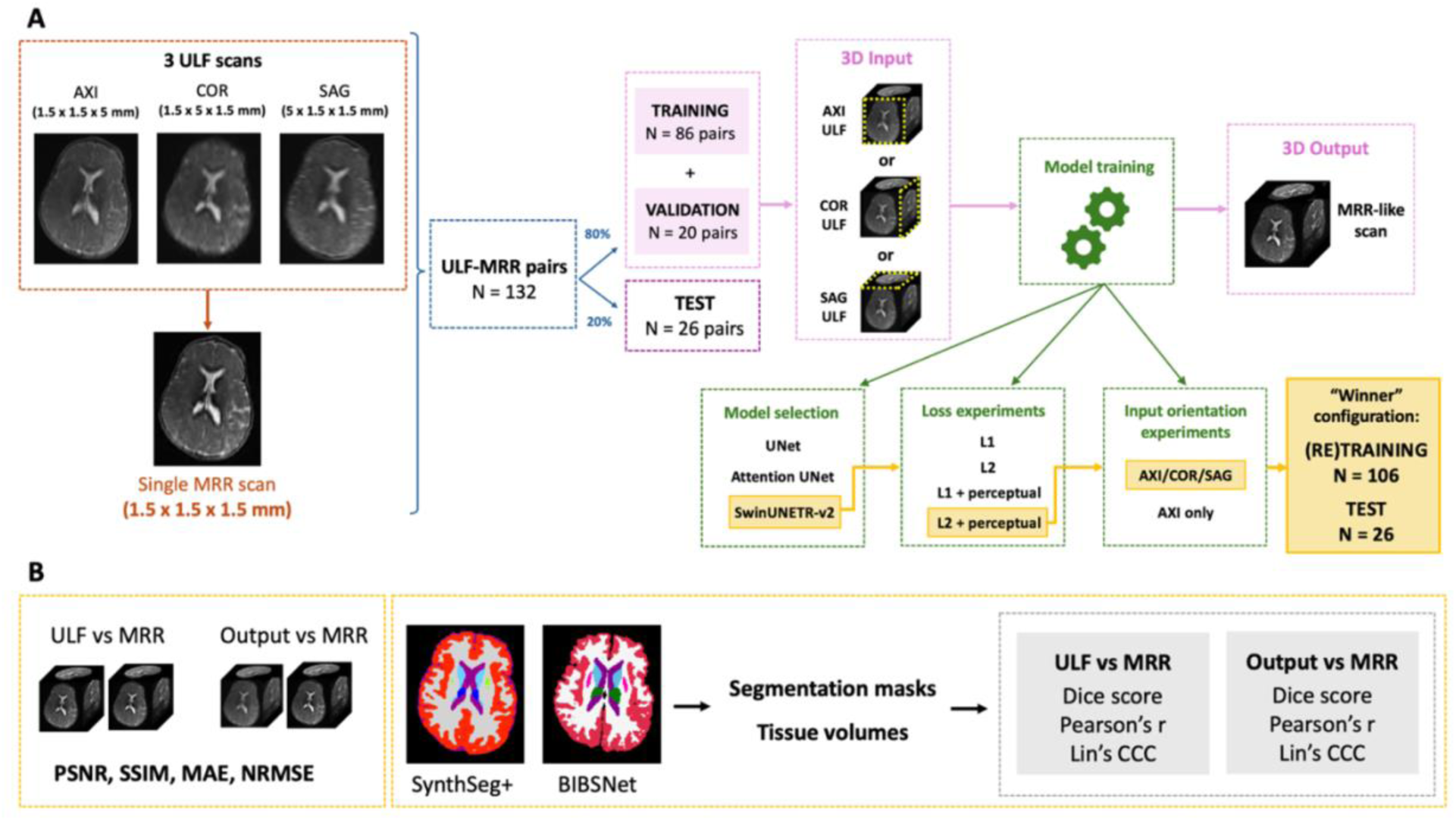
Methods workflow. A) Model training and selection. Three ULF scans are combined into a single higher-resolution MRR scan. Pairs of ULF and MRR scans are split into training, validation and test sets. Models are trained to accept a single ULF scan and output an MRR-like scan. During training, different model architectures, loss functions and input types are used to select the best performing “winner” model that is retrained on the combined training and validation datasets. **B) Model evaluation.** Image quality metrics are computed between the original ULF and the corresponding ground truth MRR scans and between model outputs and the corresponding target MRR scans. SynthSeg+ and BIBSNet segmentations are applied to model outputs to obtain segmentation masks and tissue volume estimates. Performance is evaluated compared to ULF scans and ground truth MRR using Dice overlap of tissue labels and correlations of tissue volumes.

### Model Selection, Training and Inference

To identify the best deep-learning approach for ULF super-resolution, we compared three architectures for mapping randomly sampled single ULF inputs to corresponding MRR targets: a 3D UNet [11], an Attention UNet [17], and a SwinUNETR-v2 transformer [18]. Models were trained to accept a single ULF scan as input and generate a super-resolved scan as the output, targeting the quality of a MRR scan. Training was done on 86 and validated on 20 subjects for 300 epochs using 128^3^ patches (due to GPU capacity restrictions) with extensive 3D augmentation (affine, elastic, flips, bias-field). Architecture selection used L1 loss and four evaluation metrics (peak signal-to-noise ratio - PSNR, structural similarity index - SSIM, mean absolute error - MAE and normalised root mean squared error - NRMSE) (Table S1). The best performing model across all four metrics was further optimised by testing a range of losses (L1, L2, L1+perceptual, L2+perceptual). To compute the perceptual loss for 3D MRI data, we utilised the *PerceptualLoss* module from the MONAI framework. Features were extracted using a pretrained VGG network. To accommodate the 3D nature of the MRI volumes within a 2D network architecture, the 3D inputs were processed as a stack of 2D slices. The perceptual loss was incorporated into the overall training objective alongside the standard pixel-wise loss (L1 or L2). The total loss function was defined as a weighted sum of pixel-wise loss and perceptual loss, where the perceptual loss weight was set to 0.01. In addition to optimising the loss function, we compared training on a single scan orientation (AXI) versus randomly sampled mixed orientations (AXI, COR or SAG). When training using a single orientation, the model always received an axial scan as input, resulting in a training sample of 86 scans. When training on mixed orientations, for each participant, one of the three ULF scans was randomly sampled as an input in each epoch, resulting in a more heterogenous but larger training sample of 258 scans. The corresponding MRR scan served as the target image in both the single orientation and mixed orientation cases.

Rank aggregation across all metrics determined the optimal configuration (Table S2). The best model was then retrained on the combined 106-subject training and validation set (Fig. 1A) with a learning rate of 10^-4^ using the Adam optimizer [25]. We implemented models in MONAI (PyTorch) and trained on a GPU. Using a batch size of 1 across the 106 samples, the final model underwent 1000 training epochs (approximately 90 hours on an NVIDIA Quadro RTX 8000 GPU). A larger batch size was not feasible due to GPU memory constraints.

After training, the test set was used to assess the model’s performance. This was done by taking the final weights of the model and running inference on each of the three ULF scans per test participant, to produce three super-resolved output scans for subsequent comparison to MRR ground truth scans. This resulted in 78 predictions. Inference took approximately 1 minute per participant on a modern CPU and ∼1 second per participant on a GPU.

### Model Evaluation

We assessed the performance of the best model by comparing its outputs to both the original ULF scans to quantify improvement in image quality, and to the target MRR scans to quantify how well the model learns to approximate these higher-resolution scans (Fig. 1B). In addition, we also assessed whether the quality of model outputs is influenced by the orientation of the input scan (AXI, COR or SAG).

We first evaluated the image quality using four metrics: PSNR, SSIM, MAE and NRMSE. These metrics were computed between the original ULF scans and the corresponding ground truth MRR scans and between the predicted outputs and the corresponding ground truth MRR scans. The analysis was conducted across all 26 test participants.

We used BIBSNet [26] [27] and SynthSeg+ [28] tools to segment all brain scans and estimate tissue volume for subsequent evaluation of performance using both the Dice overlap of segmented brain regions within participants and correlations of tissue volumes across participants. BIBSNet is a deep learning-based paediatric brain segmentation tool trained on high-field MRI scans of infants aged 0–8 months, enabling accurate tissue and subcortical structure segmentation in early infancy without requiring manual annotations or fine-tuning. BIBSNet produces volumetric labels for cortical and subcortical segmentations in infant brains with 29 FreeSurfer-compatible labels, making it suitable for quantitative evaluation in infant neuroimaging. Despite being trained on adult data, SynthSeg+ is a deep learning tool trained to handle brain scans acquired at any resolution or contrast, without the need for retraining or fine-tuning, making it particularly suitable for the segmentation of low-resolution 64 mT scans [29]. SynthSeg+ generates 98 labels for distinct brain regions. We performed our analyses on 4 global tissue types, namely, white matter (WM), cerebrospinal fluid (CSF), cortical grey matter (GM_cort_) and subcortical grey matter (GM_subcort_) by aggregating multiple labels. In addition to the 4 global tissue types, we repeated the analyses on 8 bilateral subcortical regions: thalamus, amygdala, accumbens, hippocampus, putamen, pallidum, caudate and ventral diencephalon. Example BIBSNet and SynthSeg+ segmentation masks for one test participant can be seen in Fig. 2. Before conducting the analyses, we performed visual QC of segmentations and excluded 4 participants with failed SynthSeg+ segmentations from further analyses (see Supplementary Fig. S2 for examples of excluded participants). SynthSeg+ also outputs automated QC scores for three global tissue types (WM, CSF, GM_cort_) and a subset of subcortical regions (thalamus, putamen + pallidum, hippocampus + amygdala). As SynthSeg+ was trained on adult scans, we did not exclude segmentations based on automated QC scores. Still, we calculated average QC scores for each region and scan type across the 22 participants with segmentations which were visually of high quality (see Supplementary Table S3).

**Figure 2:**
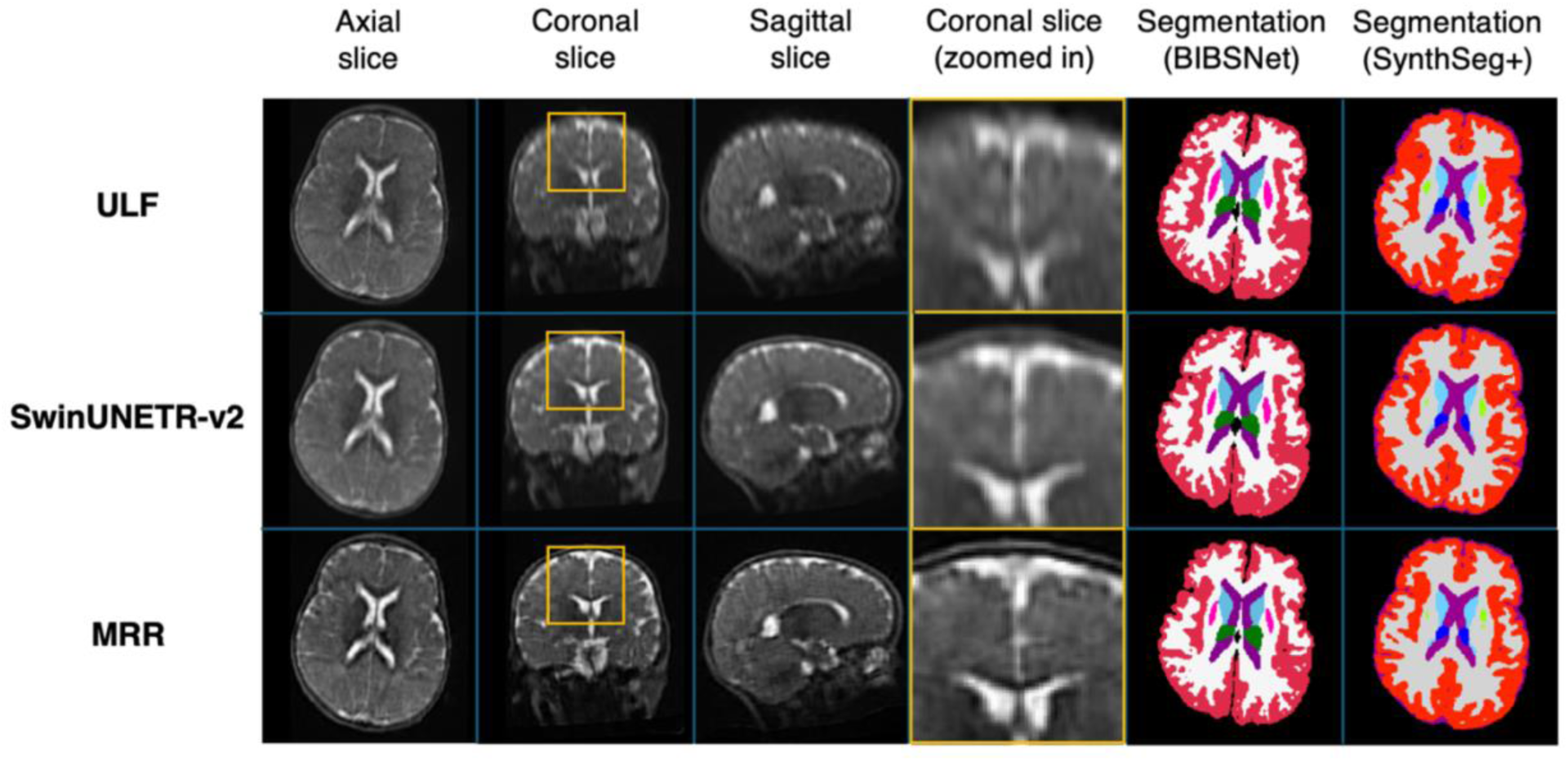
Scans from a single example test subject. The first row displays an ULF (AXI) input scan, the middle row shows the model output, and the bottom row shows the MRR scan, which serves as both the super-resolution target and reference standard. Left to right: Axial slice, coronal slice, sagittal slice, magnified features from a coronal slice, BIBSNet segmentation masks on axial slice, and SynthSeg+ segmentation masks on axial slice.

Finally, to evaluate out-of-site generalisability, we ran inference on a small sample of participants from an external ULF dataset acquired at a different scanner site (MINE dataset) [30]. Restricting the dataset to participants of the same age as the training set (i.e., 6 months) and with high-quality scans available in all three orientations (required to reconstruct MRR scans for performance evaluation), meant that only five MINE participants were eligible for inclusion in this exploratory analysis. Image quality metrics and Dice scores were computed using the same procedures as above to enable a consistent comparison.

Before applying segmentations, each participant had their original ULF scans and the three model predictions rigid-registered to their MRR scan using FSL FLIRT, a tool for linear brain image registration [31]. This was done to ensure the segmentations being compared are perfectly aligned, which is necessary for the Dice score to be meaningful.

### Statistical Analyses

Given the small sample size (*n* = 26 or *n* = 22 for individual scan orientations used for BIBSNet and SynthSeg+ respectively) and given its robustness to outliers and data distributions, we used a non-parametric Wilcoxon signed-rank test [32] to quantify the improvement in image quality following super-resolution, by comparing the Dice overlap between segmentations of ULF and MRR scans, to Dice overlap between segmentations of predicted and MRR scans. We assessed tissue volume correlations across participants using Pearson’s r [33] to measure the strength of linear correlation, as well as Lin’s concordance correlation coefficient (CCC) [34] to quantify exact agreement (i.e., alignment with the identity line, y=x).

## Results

The outputs of the SwinUNETR-v2 model are visually superior and show increased sharpness compared to the original ULF scans (Fig. 2). Model predictions show the most prominent enhancement in quality along the two low-resolution orthogonal planes, resulting in a sharper image with increased detail as seen in the enlarged portion of the brain in Fig. 2. These visual differences are also reflected in higher PSNR and SSIM and lower MAE and NRMSE values for model outputs compared to ULF input scans (Table 1).

**Table 1:** Image quality metrics for input ULF and model output scans. Peak signal-to-noise ratio (PSNR), structural similarity index measure (SSIM), mean absolute error (MAE) and normalised root-mean-squared error (NRMSE) were calculated by comparing model inputs and outputs to reference MRR scans. Values

| Comparison | PSNR (↑) | SSIM (↑) | MAE (↓) | NRMSE (↓) |
| --- | --- | --- | --- | --- |
| ULF vs MRR | $25.47 \pm 1.70$ | $0.64 \pm 0.13$ | $0.035 \pm 0.013$ | $0.054 \pm 0.011$ |
| Model vs MRR | **$28.60 \pm 1.49$** | **$0.79 \pm 0.052$** | **$0.018 \pm 0.0028$** | **$0.038 \pm 0.0066$** |

We quantified improvement in BIBSNet tissue segmentation from SR by comparing the Dice overlap of model predictions and MRR scans to the Dice overlap of ULF and MRR scans. We tested the significance of differences in individual overlap using the Wilcoxon signed -rank test (WSR) with Bonferroni correction for multiple comparisons. For CSF, WM and GM_cort_, the median Dice score increased significantly: from 0.75 to 0.76 for CSF, 0.77 to 0.78 for WM and 0.71 to 0.73 for GM_cort_ (Fig. 3 and Table 2). Dice scores did not increase significantly for GM_subcort_. Bonferroni correction for multiple comparisons was applied (i.e. *p* < 0.05/4 = 0.0125). Within individual orientations, the most noticeable improvement in SR outputs was observed for GM_cort_ in AXI and SAG scans and for CSF in COR scans (Fig. 3 and Table 2). SAG scans showed the lowest baseline Dice score, with significant improvements in the WM and GM_cort_. Additionally, we tested whether there is a significant difference between Dice scores of scans predicted from AXI, COR or SAG orientations. For WM and GM_cort_, predictions generated using AXI or COR ULF scans showed significantly higher Dice overlap to MRR targets than predictions generated using SAG ULF scans (Table 2). We found no significant difference between Dice scores of predictions generated using AXI and COR ULF scans (Table 2).

**Figure 3:**
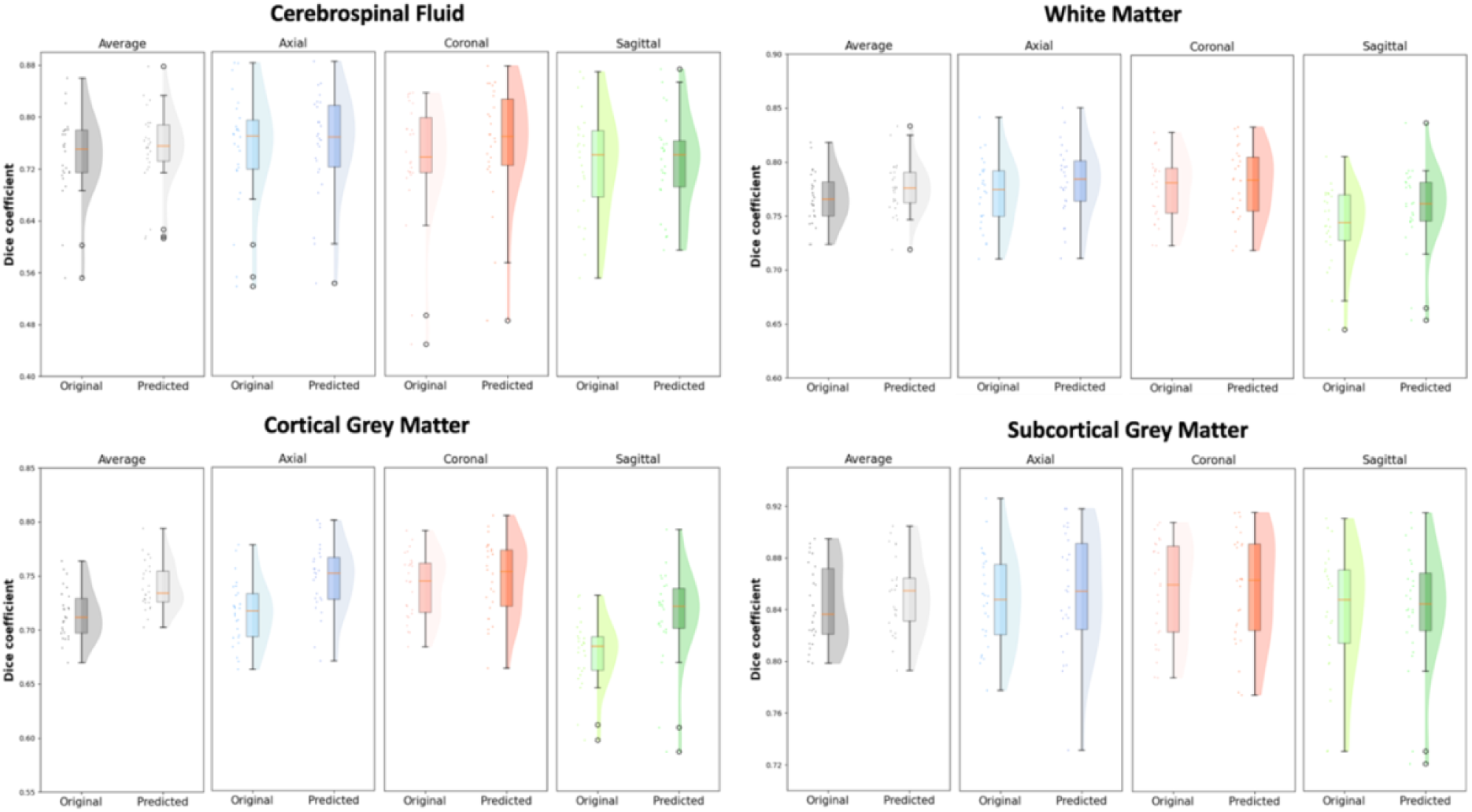
BIBSNet Dice overlap scores between model predictions and MRR scans, and between ULF and MRR scans, for 4 global tissue types. The ‘All orientations’ plots represent Dice scores obtained by combining axial, coronal and sagittal scans and have 78 data points. ‘Axial’, ‘Coronal’ and ‘Sagittal’ plots represent Dice scores obtained from each orientation separately and have 26 data points each. Original = original ULF scan, Predicted = model prediction.

**Table 2:** Wilcoxon signed-rank test applied to Dice scores obtained with BIBSNet. Wilcoxon signed-rank test applied to Dice scores between original and predicted scans (all orientations combined and across AXI, COR, and SAG), as well as between predicted Dice scores for each pair of orientations. p-value and RBC are reported for each comparison. RBC is a measure of effect size (strength of the difference between two groups) ranging between -1 and 1, where the closer the RBC is to ±1, the stronger the effect. Abbreviations: RBC = rank biserial correlation, orig. = original, pred. = predicted, CSF = cerebrospinal fluid, WM = white matter, GM_cort_ = cortical grey matter, GM_subcort_ = subcortical grey matter.

| Region | All<sub>orig</sub> vs All<sub>pred</sub><br>(orig. < pred.) |  | AXI<sub>orig</sub> vs AXI<sub>pred</sub> |  | COR<sub>orig</sub> vs COR<sub>pred</sub> |  | SAG<sub>orig</sub> vs SAG<sub>pred</sub> |  | AXI<sub>pred</sub> vs COR<sub>pred</sub> |  | AXI<sub>pred</sub> vs SAG<sub>pred</sub> |  | COR<sub>pred</sub> vs SAG<sub>pred</sub> |  |
| --- | --- | --- | --- | --- | --- | --- | --- | --- | --- | --- | --- | --- | --- | --- |
|  | p-value | RBC | p-value | RBC | p-value | RBC | p-value | RBC | p-value | RBC | p-value | RBC | p-value | RBC |
| CSF | .0088* | 0.53 | .016 | 0.48 | <.001* | 0.7 | .42 | 0.05 | .54 | -0.02 | 0.03 | 0.42 | .083 | 0.32 |
| WM | <.001* | 0.91 | <.001* | 0.98 | .0023* | 0.62 | <.001* | 0.95 | .44 | 0.04 | .016 | 0.48 | .0082* | 0.53 |
| GM<sub>cort</sub> | <.001* | 1.0 | <.001* | 0.99 | .014 | 0.49 | <.001* | 0.98 | .57 | -0.04 | .0021* | 0.62 | .0023* | 0.62 |
| GM<sub>subcort</sub> | .028 | 0.43 | .065 | 0.34 | .27 | 0.15 | .29 | 0.13 | .65 | -0.08 | .16 | 0.23 | .19 | 0.2 |
| Thalamus | .047 | 0.38 | .18 | 0.21 | .34 | 0.1 | .27 | -0.14 | .36 | 0.08 | .35 | 0.09 | .31 | 0.12 |
| Amygdala | .20 | 0.20 | .99 | -0.5 | .55 | -0.03 | .07 | 0.33 | .80 | -0.19 | .77 | -0.17 | .38 | 0.07 |
| Accumbens | .74 | -0.14 | .90 | -0.29 | .5 | 0.00 | .77 | -0.17 | .89 | -0.27 | .18 | 0.21 | .055 | 0.36 |
| Hippocampus | .21 | 0.19 | .99 | -0.52 | .23 | 0.17 | .0031* | 0.60 | .82 | -0.02 | .72 | -0.13 | .12 | 0.26 |
| Putamen | .056 | 0.36 | .10 | 0.29 | .30 | 0.12 | .39 | 0.07 | .42 | 0.05 | .032 | 0.42 | .018 | 0.47 |
| Pallidum | .38 | 0.07 | .14 | 0.25 | .43 | 0.04 | .67 | -0.10 | .59 | -0.05 | .061 | 0.35 | .020 | 0.46 |
| Caudate | .035 | 0.41 | <.001* | 0.67 | .61 | -0.06 | .62 | -0.07 | .57 | -0.04 | .040 | 0.4 | .030 | 0.42 |
| Diencephalon | .38 | 0.07 | .90 | -0.3 | .53 | -0.01 | .54 | -0.02 | .63 | -0.07 | .36 | 0.08 | .21 | 0.19 |

We performed the same analyses on 8 bilateral subcortical brain regions and here too Bonferroni correction for multiple comparisons was used (i.e. *p* < 0.05/8 = 0.00625). Considering all orientations together, all regions except the hippocampus showed an increase in Dice overlap; however, this was not significant (Table 2 and Fig. S3). When examining individual orientations, the most significant improvement if Dice overlap following SR was observed in the caudate for AXI scans and hippocampus for SAG scans. Interestingly, Dice overlap did not improve significantly for COR scans in any of the subcortical brain regions.

We further inspected correlations of tissue volume across participants, quantifying improvements in image quality following SR by comparing correlations between model outputs and MRR targets, to correlations between ULF inputs and MRR. Within the 4 global tissue types, across all scan orientations combined, we observed the greatest increases in correlation in GM_cort_, including both linear correlation (Pearson’s r rose from 0.89 to 0.98) and exact agreement (CCC rose from 0.51 to 0.95) (Fig. 4). CSF, GM_subcort_ and WM already showed high linear correlation and agreement at baseline (before SR), and therefore also smaller differences in both measures following SR (Fig. 4). When examining correlations within each specific orientation separately, a less obvious improvement was observed (Table 3). AXI scans showed an increase in volume correlations only for GM_cort_, COR scans only for WM and GM_subcort_ and SAG scans showed an increase in volume correlations only for CSF and GM_cort_ (Table 3).

**Figure 4:**
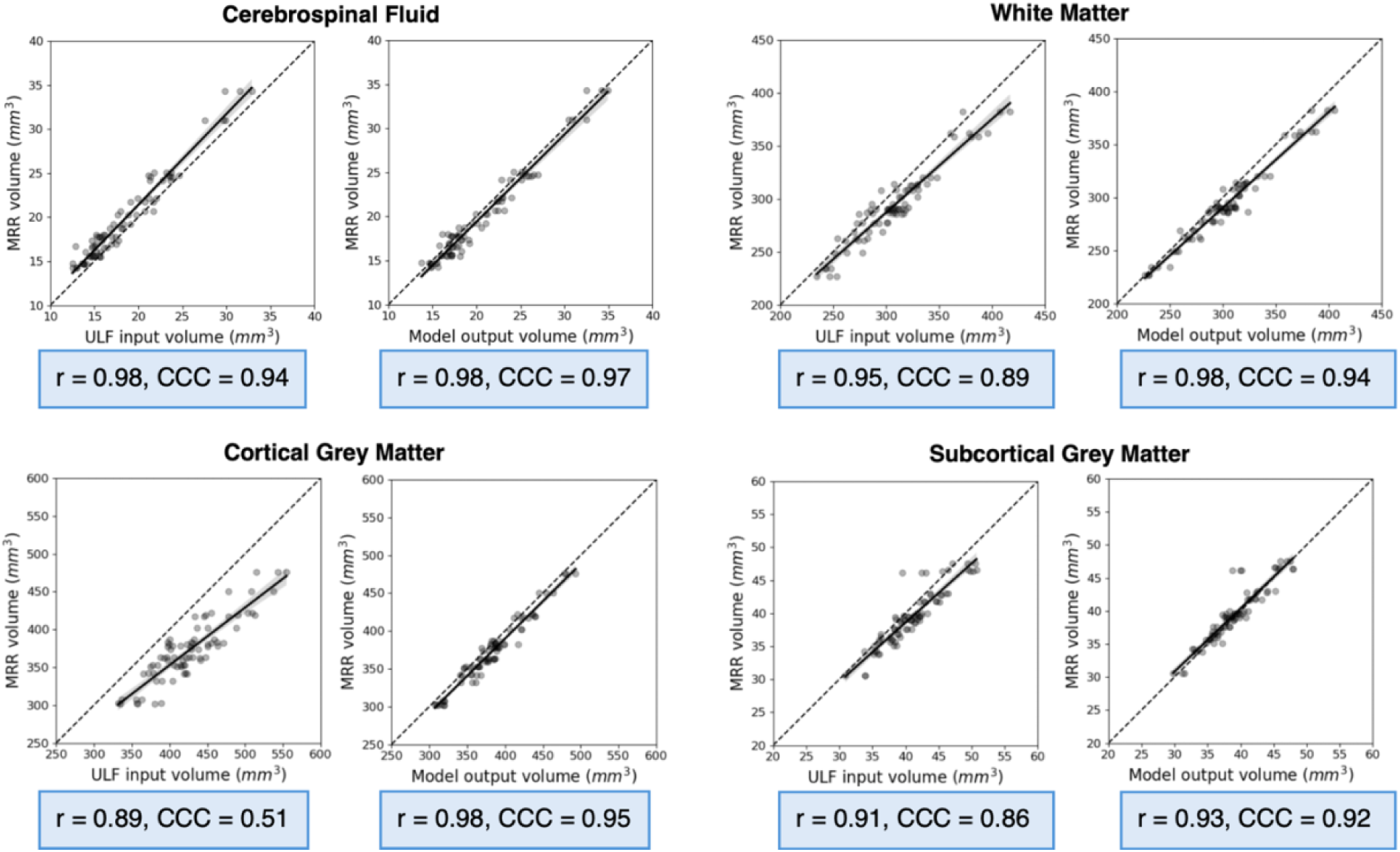
Tissue volume correlations between MRR and ULF versus model output scans for 4 global tissue types obtained with BIBSNet. Blue boxes under the correlation plots report Pearson’s correlation coefficient and Lin’s concordance correlation coefficient (CCC).

**Table 3:** Volume correlations obtained with BIBSNet for 4 global tissue types per orientation. Volume correlations between MRR and ULF, MRR and model prediction volumes and differences between them (Δ Pearson’s r and Δ CCC).

| Region | AXI |  | COR |  | SAG |  |
| --- | --- | --- | --- | --- | --- | --- |
|  | Pearson's r | CCC | Pearson's r | CCC | Pearson's r | CCC |
|  | Volume correlation between MRR and ULF |  |  |  |  |  |
| CSF | 0.99 | 0.98 | 0.99 | 0.87 | 0.99 | 0.97 |
| WM | 0.99 | 0.87 | 0.94 | 0.94 | 0.98 | 0.87 |
| GM<sub>cort</sub> | 0.93 | 0.52 | 0.76 | 0.76 | 0.97 | 0.39 |
| GM<sub>subcort</sub> | 0.94 | 0.83 | 0.91 | 0.91 | 0.95 | 0.85 |
| Volume correlation between MRR and model prediction |  |  |  |  |  |  |
| CSF | 0.98 | 0.96 | 0.98 | 0.98 | 0.99 | 0.98 |
| WM | 0.98 | 0.94 | 0.98 | 0.97 | 0.98 | 0.91 |
| GM<sub>cort</sub> | 0.97 | 0.93 | 0.97 | 0.97 | 0.99 | 0.95 |
| GM<sub>subcort</sub> | 0.93 | 0.93 | 0.92 | 0.91 | 0.94 | 0.94 |
| $\Delta$ volume correlation (MRR vs model prediction - MRR vs ULF) | | | | | | |
| CSF | -0.01 | -0.02 | -0.01 | 0.11 | 0 | 0.01 |
| WM | -0.01 | 0.07 | 0.04 | 0.03 | 0 | 0.04 |
| GM<sub>cort</sub> | 0.04 | 0.41 | 0.21 | 0.21 | 0.02 | 0.56 |
| GM<sub>subcort</sub> | -0.01 | 0.10 | 0.01 | 0.0 | -0.01 | 0.09 |

We next inspected differences in volume correlations following SR in individual (bilateral) subcortical regions. Across the three orientations combined, improvements were observed for all subcortical regions except for the thalamus (Fig. S4), with the greatest improvements in the amygdala, accumbens and ventral diencephalon (Pearson’s r rose from 0.81 to 0.87, from 0.75 to 0.88 and from 0.80 to 0.84, respectively) (Fig. S4). Within individual orientations, AXI scans showed improvements in linear correlation for the hippocampus, putamen, pallidum and ventral diencephalon and improvements in exact agreement for all subcortical regions except the accumbens (Table 4). SAG scans showed improvements in linear correlation for the amygdala, accumbens, pallidum and caudate (Table 4). We replicated the main trends in results with SynthSeg+, which showed greater improvements in volume correlations and Dice overlap following super-resolution (see Supplementary Tables S4-6 and Supplementary Figures S5-8).

**Table 4:** Volume correlations obtained with BIBSNet for 8 subcortical regions per orientation. Volume correlations between MRR and ULF, MRR and model predictions and differences between them (Δ Pearson’s r and Δ CCC).

| Region | AXI |  | COR |  | SAG |  |
| --- | --- | --- | --- | --- | --- | --- |
|  | Pearson's r | CCC | Pearson's r | CCC | Pearson's r | CCC |
| Volume correlation between MRR and ULF |  |  |  |  |  |  |
| Thalamus | 0.89 | 0.78 | 0.86 | 0.82 | 0.89 | 0.65 |
| Amygdala | 0.94 | 0.67 | 0.91 | 0.90 | 0.79 | 0.59 |
| Accumbens | 0.87 | 0.87 | 0.92 | 0.87 | 0.76 | 0.47 |
| Hippocampus | 0.91 | 0.72 | 0.87 | 0.86 | 0.94 | 0.88 |
| Putamen | 0.91 | 0.83 | 0.93 | 0.93 | 0.88 | 0.87 |
| Pallidum | 0.87 | 0.83 | 0.86 | 0.82 | 0.80 | 0.79 |
| Caudate | 0.94 | 0.87 | 0.93 | 0.93 | 0.87 | 0.86 |
| Diencephalon | 0.83 | 0.81 | 0.90 | 0.85 | 0.84 | 0.71 |
| Volume correlation between MRR and model prediction |  |  |  |  |  |  |
| Thalamus | 0.84 | 0.83 | 0.86 | 0.84 | 0.86 | 0.80 |
| Amygdala | 0.91 | 0.87 | 0.90 | 0.88 | 0.84 | 0.83 |
| Accumbens | 0.87 | 0.86 | 0.94 | 0.93 | 0.88 | 0.83 |
| Hippocampus | 0.94 | 0.93 | 0.88 | 0.87 | 0.93 | 0.86 |
| Putamen | 0.92 | 0.92 | 0.93 | 0.88 | 0.87 | 0.87 |
| Pallidum | 0.90 | 0.89 | 0.87 | 0.73 | 0.84 | 0.67 |
| Caudate | 0.92 | 0.90 | 0.92 | 0.88 | 0.88 | 0.80 |
| Diencephalon | 0.85 | 0.83 | 0.87 | 0.73 | 0.83 | 0.78 |
| $\Delta$ volume correlation (MRR vs model prediction - MRR vs ULF) | | | | | | |
| Thalamus | -0.05 | 0.02 | 0.0 | 0.02 | -0.03 | 0.15 |
| Amygdala | -0.03 | 0.2 | -0.01 | -0.02 | 0.05 | 0.24 |
| Accumbens | 0.0 | -0.01 | 0.03 | 0.06 | 0.12 | 0.36 |
| Hippocampus | 0.3 | 0.21 | 0.01 | 0.01 | -0.01 | -0.02 |
| Putamen | 0.1 | 0.09 | 0.0 | 0.05 | -0.01 | 0.0 |
| Pallidum | 0.03 | 0.06 | 0.01 | -0.09 | 0.04 | -0.12 |
| Caudate | -0.02 | 0.03 | -0.01 | -0.05 | 0.01 | -0.06 |
| Diencephalon | 0.02 | 0.02 | -0.03 | -0.12 | -0.01 | 0.07 |

On the out-of-site MINE dataset, improvements in image quality and Dice overlap were not consistently observed (Supplementary Tables S7–S8).

## Discussion

ULF MRI scanners offer a more accessible and affordable alternative for neuroimaging in LMICs, but come at the cost of reduced image quality. This study explores whether deep learning, specifically a SwinUNETR-v2 model, can enhance the resolution of individual ULF scans to match MRR scans, which traditionally require three orthogonal acquisitions. The results show that the model significantly improved the quality of ULF scans, both globally and locally. Dice overlap scores for global tissue types and most subcortical regions were significantly higher in model outputs compared to ULF inputs. Additionally, volume correlations between predicted and MRR scans were stronger for all global tissue types except CSF and for most individual deep-brain structures, compared to volume correlations between original ULF and MRR scans. This has been demonstrated using two different segmentations tools. Overall, our findings support the effectiveness of deep learning enhancement of individual ULF scans without paired HF scans, resulting in a closer approximation of MRR-quality outputs.

The ability to reconstruct a higher-resolution scan from any single ULF scan (axial, coronal or sagittal) is an improvement to existing SR approaches, that either require all three ULF scans to reconstruct a higher-resolution scan [19], or require paired ULF and HF data for model training [14–16]. Our approach has multiple advantages. Firstly, the ability to generate higher-quality scans from single ULF acquisitions could reduce scanning time, making MRI data acquisition more suitable for scanning infants who cannot keep still for long periods of time and often struggle with extended scanning sessions [35]. Shorter scanning sessions are also cheaper, making this approach accessible for adoption in LMICs, which could further boost neurodevelopmental research and establish the technical feasibility required for future clinical applications. Moreover, since the model requires only a single ULF scan, our approach could save significant amounts of data in instances where participants may be excluded from studies due to not having high-quality scans from all three orientations.

While our approach requires high-quality ULF data from all three orientations to train a super-resolution model, it does not require paired HF scans. One advantage of this approach is that our model can be easily retrained at individual ULF scanner sites without access to HF MRI, on age- and/or diagnosis-specific populations. The other important benefit is that we avoid the domain gap between input and target data observed in several other super-resolution studies where models are trained on paired ULF and HF scans, or even HF scans paired with synthetically down-sampled HF counterparts, which may not generalise well to empirical ULF scans [14] [15] [36]. The domain gap is problematic as morphometric measurements are highly dependent on the MR image contrast and noise, both of which are influenced by the field strength of the MR system [29] [37]. Additionally, the domain gap presents previously used SR models with an additional translation task that they have to learn. Alongside translation from low-resolution to higher-resolution and from low SNR to high SNR, these models have to learn to bridge the domain gap by mapping between scans of subtly different contrast that depends on field strength. Our model is not presented with such a challenge and therefore has a slightly easier task to learn. It is worth acknowledging, however, that models trained on paired ULF and HF scans can learn to output higher-quality images compared to ours.

Still, our model was presented with other challenges. The contrast between grey and white matter is the least distinct at 6 months of age due to active myelination processes [21] [22] [38], making the learning particularly demanding. Despite this, the model consistently improved ULF image quality and produced outputs that closely approximated the MRR targets. Additionally, to enable the super-resolution of ULF scans acquired in any orientation, the model had to learn mappings from three different ULF orientations (AXI, COR, SAG) to the MRR target, arguably making the task more difficult compared to learning from a single input orientation. While we also evaluated training on a single orientation (only AXI), all models were tested on all three orientations, which requires the model to generalise across inputs. Training with multiple orientations provided a three-fold increase in available training data and overall yielded better performance than single-orientation training, suggesting that the model benefits from the increased variability and data volume.

We found that when looking at global and subcortical tissue types, among the three orientations, median Dice scores obtained from BIBSNet segmentations were the highest in axial and coronal scans for both ULF and model output scans. This suggests that in situations where only a single ULF anisotropic scan can be acquired, acquisition of an axial or coronal scan should potentially be prioritised. Some differences between orientations were also observed in the study by Váša and colleagues [29], where T2-weighted ULF COR scans showed the highest correspondence to scans acquired with a 3 T HF scanner, followed by AXI and SAG scans. It is important to note, however, that this latter study focused on adult participants.

The complexity of working with infant brain data extends beyond resolution enhancement, particularly with respect to accurate segmentation. In this study, scans were segmented using two state-of-the-art automated tools: SynthSeg+ [28], a widely used method trained on adult MRI data and designed to operate across a broad range of contrasts and resolutions, and BIBSNet, a deep learning tool trained on paediatric MRI data spanning early infancy [26]. Infant brains exhibit markedly reduced grey-white matter contrast compared to adults [38], which poses challenges for segmentation, yet BIBSNet benefits from training on age-appropriate anatomy, while SynthSeg+ supports segmentation across imaging resolutions, including ULF scans. Consistent trends were observed across both tools in tissue volume correlations, while Dice scores showed improvements following super-resolution for both, but of differing magnitudes. Taken together, these results highlight both the difficulty of segmenting infant ULF brain scans and the benefit of using more than one segmentation tool for evaluation. Continued development of infant-specific approaches, such as the emerging ULF paediatric segmentation tool MiniMORPH [39], will be important for improving quantitative analyses in early neurodevelopment.

An important limitation of this study is that the data used to train the models was acquired from a single scanning site in Cape Town, from neurologically healthy 6-month-old infants only. Having such a uniform sample likely limits the generalisability of the model, especially because age can introduce significant anatomical variations that are not captured by our training set [40]. However, it is important to note this cohort is from a context where there are high instances of psychosocial adversity and disease burden, including exposure to viruses such as HIV and substances such as alcohol during pregnancy. These adversities may contribute to greater variability in brain development within the cohort [41], potentially enhancing the diversity of the training data. To assess cross-site generalisability, we conducted a small exploratory external validation on five 6-month-old infants from the independent MINE dataset [30], which showed modest and inconsistent improvements in performance. These inconsistencies are likely caused by domain shifts between datasets, such as differences in scanner software versions (e.g., versions 8.2.0–8.6.1 used for the training set versus 8.5.1–8.8.1 for the external test set), associated minor variations in acquisition protocols (e.g., varying echo times across scans), or varying environmental noise profiles related to local power supplies and electrical interference. Given the small sample size and therefore exploratory nature of this analysis, results need to be interpreted with caution. Consequently, until these analyses can be repeated using a larger external test set, out-of-site deployment of our approach would require site-specific retraining. Nonetheless, because our approach does not require paired ULF and HF data, but only relies on three ULF scans, retraining on new populations and scanner sites remains feasible, including in LMIC settings. Clinical validation of the super-resolved outputs was beyond the scope of the present work and remains an important direction for future studies.

Finally, while we show that the model can successfully super-resolve brain MRI scans, future research could investigate other deep learning architectures for the super-resolution of single ULF MRI scans without paired HF data. Such architectures include, but are not limited to, diffusion models, which have already been proposed as a paradigm for medical image reconstruction and enhancement [42] [43], and state-space models, which have shown great promise in MRI SR tasks [44].

In conclusion, the current study demonstrates the efficacy of deep learning in enhancing the resolution of ULF infant MRI scans, significantly improving tissue volume estimation and segmentation accuracy compared to original scans. By enabling the reconstruction of higher resolution brain scans from a single ULF scan without paired HF data, the model not only reduces the scanning time – which is critical for paediatric imaging – but also paves the way for global adoption of MRI technology in LMICs, potentially advancing neurodevelopmental research in these regions.

## Funding Acknowledgments

This study was supported by the Bill and Melinda Gates Foundation UNITY project [INV-032788; INV-047888; INV-004939], the Wellcome Leap 1kD programme (The First 1000 Days) [222076/Z/20/Z] and by the DELTAS II Africa Programme(Del-22-002) which is funded by the Science for Africa Foundation with support from the Wellcome Trust and the UK Foreign, Commonwealth and Development Office and is part of the the EDCTP 2 programme supported by the European Union.

## Khula South Africa Study Team

Donna Herr^2^, Marlie Miles^2^, Chloë A. Jacobs^2^, Sadeeka Williams^2^, Zamazimba Madi^2^, Nwabisa Mlandu^2^, Tembeka Mhlakwaphalwa^2^, Lauren Davel^2^, Reese Samuels^2^, Zayaan Goolam^2^, Thandeka Mazubane^2^, Bokang Methola^2^, Khanyisa Nkubungu^2^, Candice Knipe^2^ & Tracy Pan^2^

## Notes

### Competing Interest Statement

The authors have declared no competing interest.

### Summary of Updates

We expanded our architectural benchmarking to compare multiple super-resolution models for comparison with state-of-the-art approaches, added ablation analyses testing alternative loss functions and training configurations, improved our segmentation-based evaluation by integrating a paediatric segmentation tool, increased the effective test sample size for segmentation analyses and conducted an exploratory out-of-site validation using an independent dataset. The manuscript structure has been comprehensively updated to reflect these analyses, encompassing revised results sections, extended supplementary materials, and the UNet architecture figure and accompanying description replaced with a methods diagram which more specifically illustrates the unique contributions of our study. In addition, we added several references to acknowledge recently-published related work. Finally, we added Hajer Karoui, Sadia Parkar, Sidra Kaleem, and Salman Osmani as co-authors to reflect their substantial contributions to this expanded analytical framework.

## References

1. Ryan, M. E., & Jaju, A. Revolutionizing pediatric neuroimaging: the era of CT, MRI, and beyond. Child’s Nerv. Syst., 39, 10; 10.1007/s00381-023-06041-9 (2023)

2. Deoni, S. C. L. et al. Accessible pediatric neuroimaging using a low field strength MRI scanner. NeuroImage, 238, 118273. 10.1016/j.neuroimage.2021.118273 (2021)

3. Anazodo, U. C., et al. A framework for advancing sustainable magnetic resonance imaging access in Africa. NMR in Biomed., 36, 3. 10.1002/nbm.4846 (2023)

4. Ogbole, G. I., Adeyomoye, A. O., Badu-Peprah, A., Mensah, Y., & Nzeh, D. A. (2018). Survey of magnetic resonance imaging availability in West Africa. PAMJ, 30, 240. 10.11604/pamj.2018.30.240.14000 (2018)

5. Jalloul, M., et al. MRI scarcity in low- and middle-income countries. NMR in Biomed., 36, 12. 10.1002/nbm.5022 (2023)

6. Wald, L. L., McDaniel, P. C., Witzel, T., Stockmann, J. P., & Cooley, C. Z. Low-cost and portable MRI. J. Magn. Reson. Imaging, 52, 3, 686–696. 10.1002/jmri.26942 (2019)

7. Arnold, T. C., Freeman, C. W., Litt, B., & Stein, J. M. (2022). Low-field MRI: Clinical Promise and Challenges. J. Magn. Reson. Imaging, 57, 1. 10.1002/jmri.28408 (2022)

8. van Beek, E. J. R. et al. Value of MRI in medicine: More than just another test? J. Magn. Reson. Imaging, 49, 7. 10.1002/jmri.26211 (2018)

9. Abate, F. et al. UNITY: A Low-Field Magnetic Resonance Neuroimaging Initiative to Characterize Neurodevelopment in Low and Middle-Income Settings. Dev. Cog. Neurosci., 69, 101397–101397. 10.1016/j.dcn.2024.101397 (2024)

10. Li, Z. et al. DeepVolume: Brain Structure and Spatial Connection-Aware Network for Brain MRI Super-Resolution. IEEE Trans. Cyber., 51, 7, 3441–3454. 10.1109/tcyb.2019.2933633 (2021)

11. Çiçek, Ö., Abdulkadir, A., Lienkamp, S. S., Brox, T., & Ronneberger, O. 3D u-net: Learning dense volumetric segmentation from sparse annotation. Preprint at 10.48550/arxiv.1606.06650 (2016)

12. Ronneberger, O., Fischer, P., & Brox, T. U-Net: Convolutional Networks for Biomedical Image Segmentation. Preprint at https://arxiv.org/abs/1505.04597 (2015)

13. Iglesias, J. E. et al. Joint super-resolution and synthesis of 1 mm isotropic MP-RAGE volumes from clinical MRI exams with scans of different orientation, resolution and contrast. NeuroImage, 237, 118206. 10.1016/j.neuroimage.2021.118206 (2021)

14. Baljer, L., et al. Ultra-Low-Field Paediatric MRI in Low- and Middle-Income Countries: Super-Resolution Using a Multi-Orientation U-Net. Hum. Brain Mapp., 46, 1. 10.1002/hbm.70112 (2025)

15. Islam, K. et al. Improving portable low-field MRI image quality through image-to-image translation using paired low- and high-field images. Sci. Rep., 13, 21183. 10.1038/s41598-023-48438-1 (2023)

16. de Leeuw den Bouter, M. L., et al. Deep learning-based single image super-resolution for low-field MR brain images. Sci. Rep., 12, 6362. 10.1038/s41598-022-10298-6 (2022)

17. Oktay, O., et al. Attention U-Net: Learning Where to Look for the Pancreas. Preprint at 10.48550/arXiv.1804.03999 (2018)

18. Hatamizadeh, A., et al. Swin UNETR: Swin Transformers for Semantic Segmentation of Brain Tumors in MRI Images. Preprint at 10.48550/arxiv.2201.01266 (2022)

19. Deoni, S. C. L., O’Muircheartaigh, J., Ljungberg, E., Huentelman, M., & Williams, S. C. R. Simultaneous high-resolution T_2_-weighted imaging and quantitative T_2_ mapping at low magnetic field strengths using a multiple TE and multi-orientation acquisition approach. Magn. Res. Med., 88, 1273–1281. 10.1002/mrm.29273 (2022)

20. Zieff, M. R. et al. Characterizing developing executive functions in the first 1000 days in South Africa and Malawi: The Khula Study. Wellcome Open Research, 9. 10.12688/wellcomeopenres.19638.1 (2024)

21. Barkovich, A. J. Magnetic resonance techniques in the assessment of myelin and myelination. J. Inh. Metab. Dis., 28, 311–343. 10.1007/s10545-005-5952-z (2005)

22. Prastawa, M., Gilmore, J. H., Lin, W., & Gerig, G. Automatic segmentation of MR images of the developing newborn brain. Med. Image Anal., 9, 457–466. 10.1016/j.media.2005.05.007 (2005)

23. Penny, W. D., Friston, K. J., Ashburner, J. T., Kiebel, S. J. K., & Nichols, T. E. Statistical Parametric Mapping: The Analysis of Functional Brain Images (1st ed.). (Academic Press, 2006)

24. Avants, B. B., Tustison, N., & Song, G. Advanced normalization tools (ANTS). Insight J., 2, 1–35. 10.54294/uvnhin (2009)

25. Kingma, D., & Ba, J. Adam: A Method for Stochastic Optimization. Preprint at 10.48550/arXiv.1412.6980 (2014)

26. Hendrickson, T. et al. BIBSNet: A Deep Learning Baby Image Brain Segmentation Network for MRI Scans. Preprint at 10.1101/2023.03.22.533696 (2023)

27. Houghton, A., et al. BIBSnet (3.4.2). *Zenodo*. 10.5281/zenodo.13743295 (2024)

28. Billot, B., et al. Robust machine learning segmentation for large-scale analysis of heterogeneous clinical brain MRI datasets. PNAS, 120. 10.1073/pnas.2216399120 (2023)

29. Váša, F., et al. Ultra-low-field brain MRI morphometry: test-retest reliability and correspondence to high-field MRI. Imaging. Neurosci., 3. 10.1162/imag.a.930 (2025)

30. Surani, Z. et al. Maternal and environmental Impact assessment on Neurodevelopment in Early childhood years (MINE): a prospective cohort study protocol from a low, middle-income country. BMJ Open, 13, 7. 10.1136/bmjopen-2022-070283 (2023)

31. Jenkinson, M., Bannister, P., Brady, M., & Smith, S. Improved Optimization for the Robust and Accurate Linear Registration and Motion Correction of Brain Images. NeuroImage, 17, 825–841. 10.1006/nimg.2002.1132 (2002)

32. Wilcoxon, F. Individual Comparisons by Ranking Methods. Biometrics Bull., 1(6), 80–83. 10.2307/3001968 (1945)

33. Pearson, K. Notes on Regression and Inheritance in the Case of Two Parents. Proc. R. Soc. Lond., 58, 240–242. 10.1098/rspl.1895.0041 (1895)

34. Lin, L. I-Kuei. A Concordance Correlation Coefficient to Evaluate Reproducibility. Biometrics, 45, 255. 10.2307/2532051 (1989).

35. Barkovich, M. J., Li, Y., Desikan, R. S., Barkovich, A. J., & Xu, D. Challenges in pediatric neuroimaging. NeuroImage, 185, 793–801. 10.1016/j.neuroimage.2018.04.044 (2019)

36. Iglesias, J. E. et al. SynthSR: A public AI tool to turn heterogeneous clinical brain scans into high-resolution T1-weighted images for 3D morphometry. Sci. Adv., 9. 10.1126/sciadv.add3607 (2023)

37. Laguna, S. et al. Super-resolution of portable low-field MRI in real scenarios: integration with denoising and domain adaptation. MIDL 2022. https://openreview.net/forum?id=pinw0Gcot4T (2022)

38. Dubois, J. et al. MRI of the Neonatal Brain: A Review of Methodological Challenges and Neuroscientific Advances. J. Magn. Res. Imaging, 53, 1318–1343. 10.1002/jmri.27192 (2020)

39. Casella, C. MiniMORPH: A Morphometry Pipeline for Low-Field MRI in Infants. Preprint at 10.1101/2025.07.01.25330469 (2025)

40. Knickmeyer, R. C. et al. A Structural MRI Study of Human Brain Development from Birth to 2 Years. J. Neurosci., 28, 12176–12182. 10.1523/jneurosci.3479-08.2008 (2008)

41. Miguel, P. M., Pereira, L. O., Silveira, P. P., & Meaney, M. J. Early environmental influences on the development of children’s brain structure and function. Dev. Med. Child Neurol., 61, 1127–1133. 10.1111/dmcn.14182 (2019)

42. Kazerouni, A. et al. Diffusion models in medical imaging: A comprehensive survey. Med. Image Anal., 88, 102846. 10.1016/j.media.2023.102846 (2023)

43. Kim, J., & Park, H. Adaptive Latent Diffusion Model for 3D Medical Image to Image Translation: Multi-modal Magnetic Resonance Imaging Study. Preprint at https://arxiv.org/abs/2311.00265 (2023)

44. Baljer, L. et al. GAMBAS: Generalised-Hilbert Mamba for Super-resolution of Paediatric Ultra-Low-Field MRI. Preprint at 10.48550/arXiv.2504.04523 (2025)

