## Supplementary Information for "Deep learning super-resolution of paediatric ultra-low-field MRI without paired high-field scans"

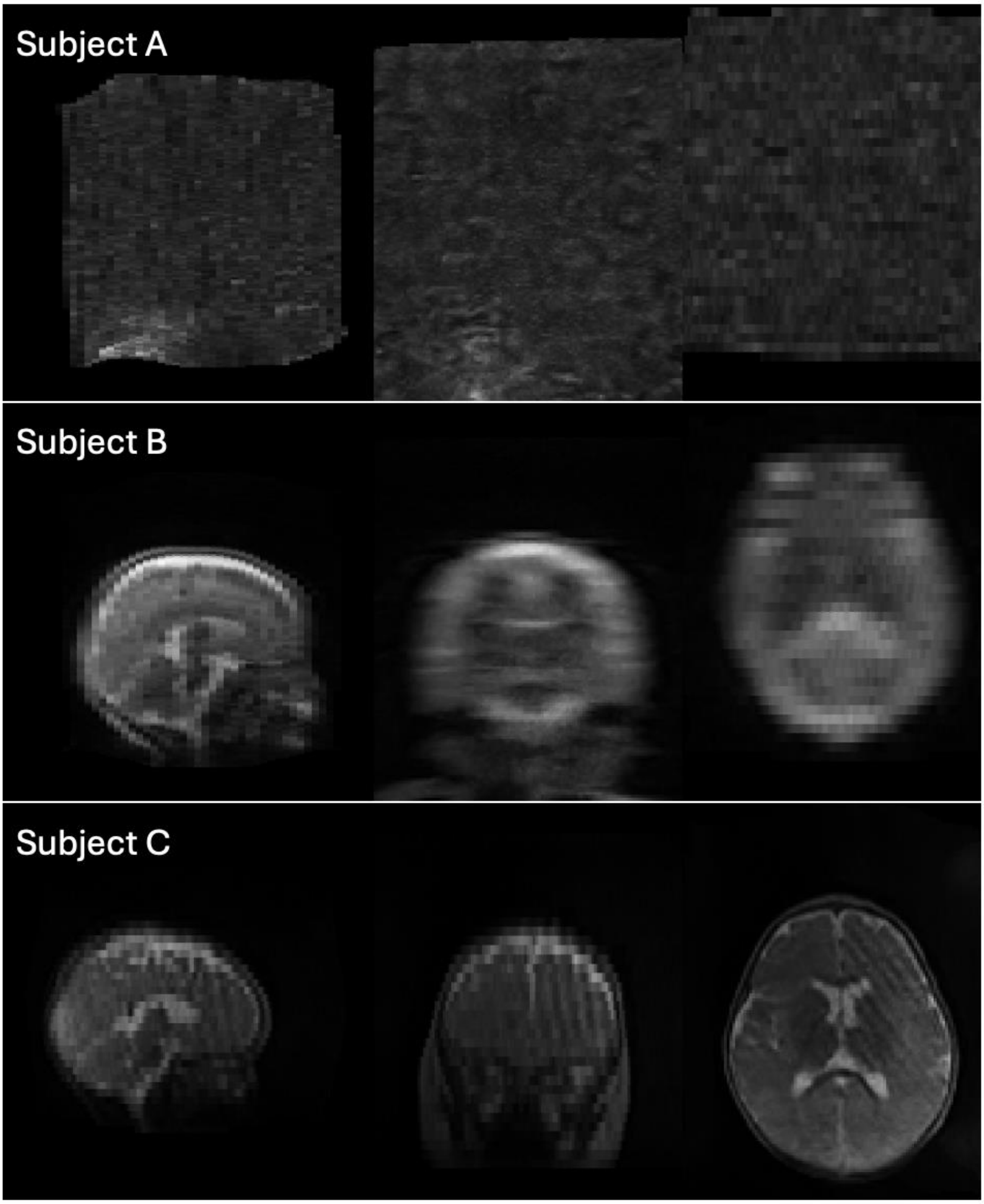


**Figure S1: Examples of subjects excluded from the study.** All excluded scans exhibited severe motion artifacts or other significant imaging artifacts.

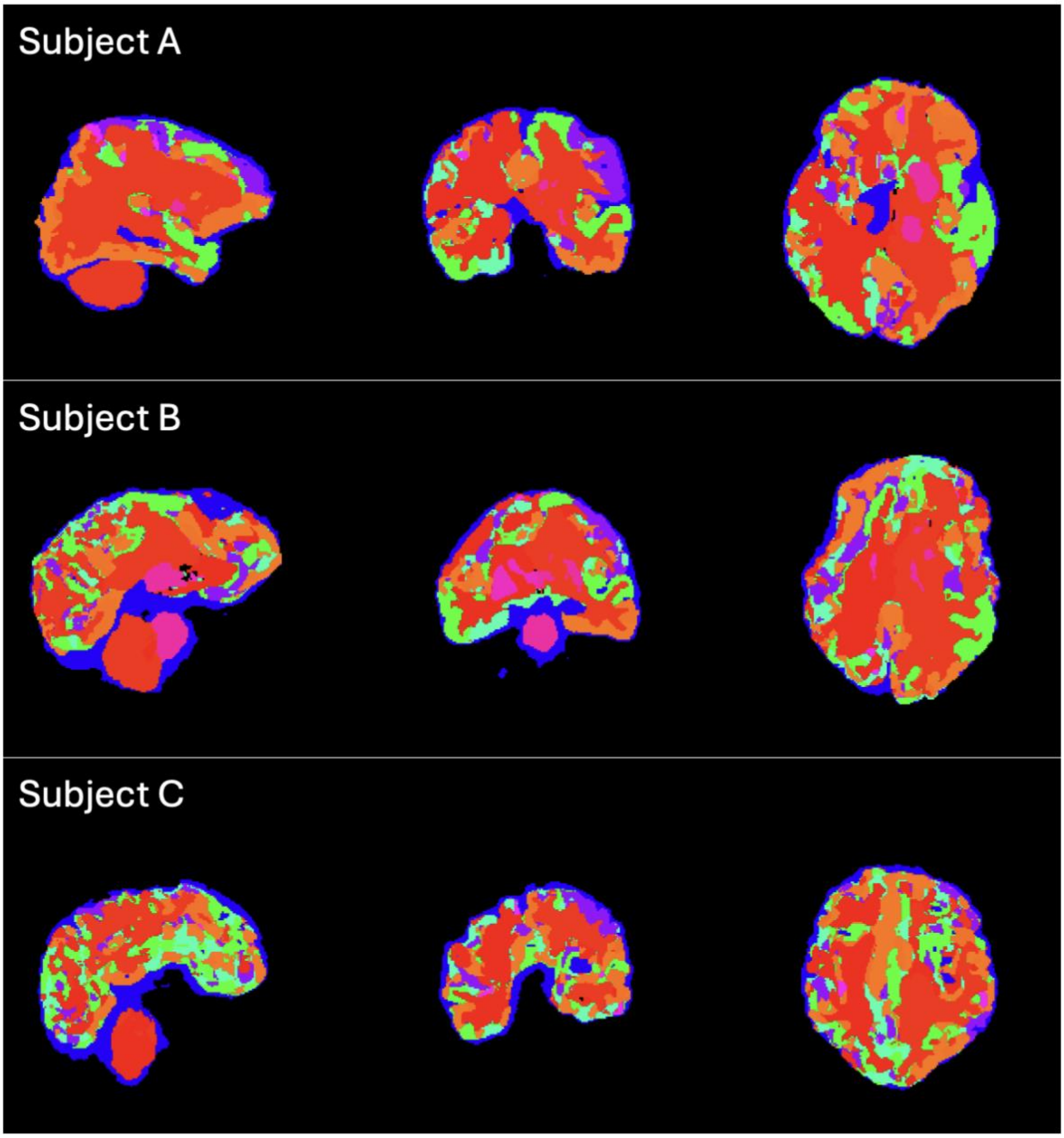


**Figure S2: Examples of excluded subjects due to failed Synthseg+ segmentations.** Segmented scans fail to accurately capture individual brain regions in all three orientations.

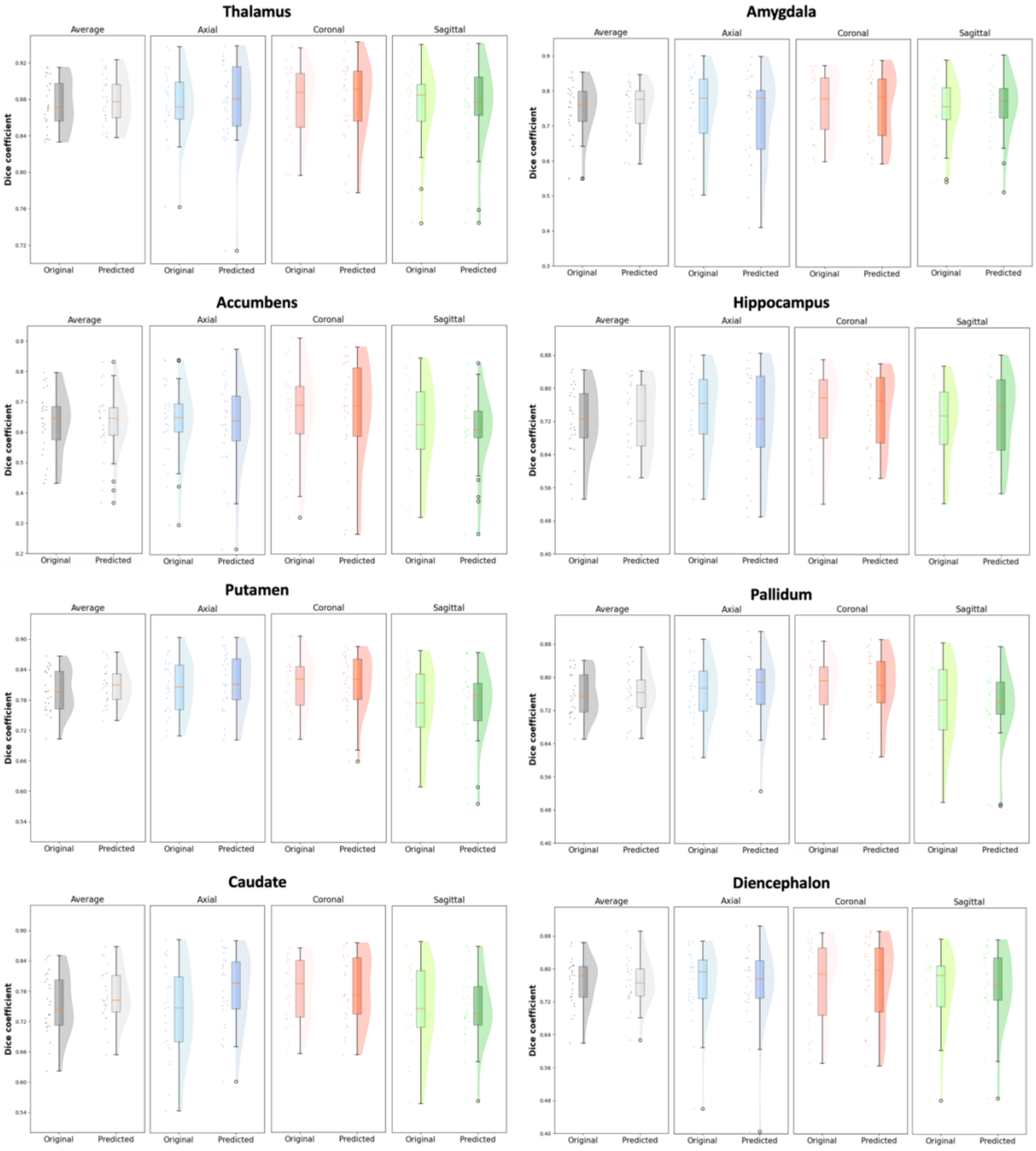


**Figure S3: BIBSNet Dice overlap scores between model predictions and MRR scans, and between ULF and MRR scans, for 8 subcortical regions.** The ‘All orientations’ plots represent Dice scores obtained by combining axial, coronal and sagittal scans and have 78 data points. ‘Axial’, ‘Coronal’ and ‘Sagittal’ plots represent Dice scores obtained from each orientation separately and have 26 data points each. Original = original ULF scan, Predicted = model prediction.

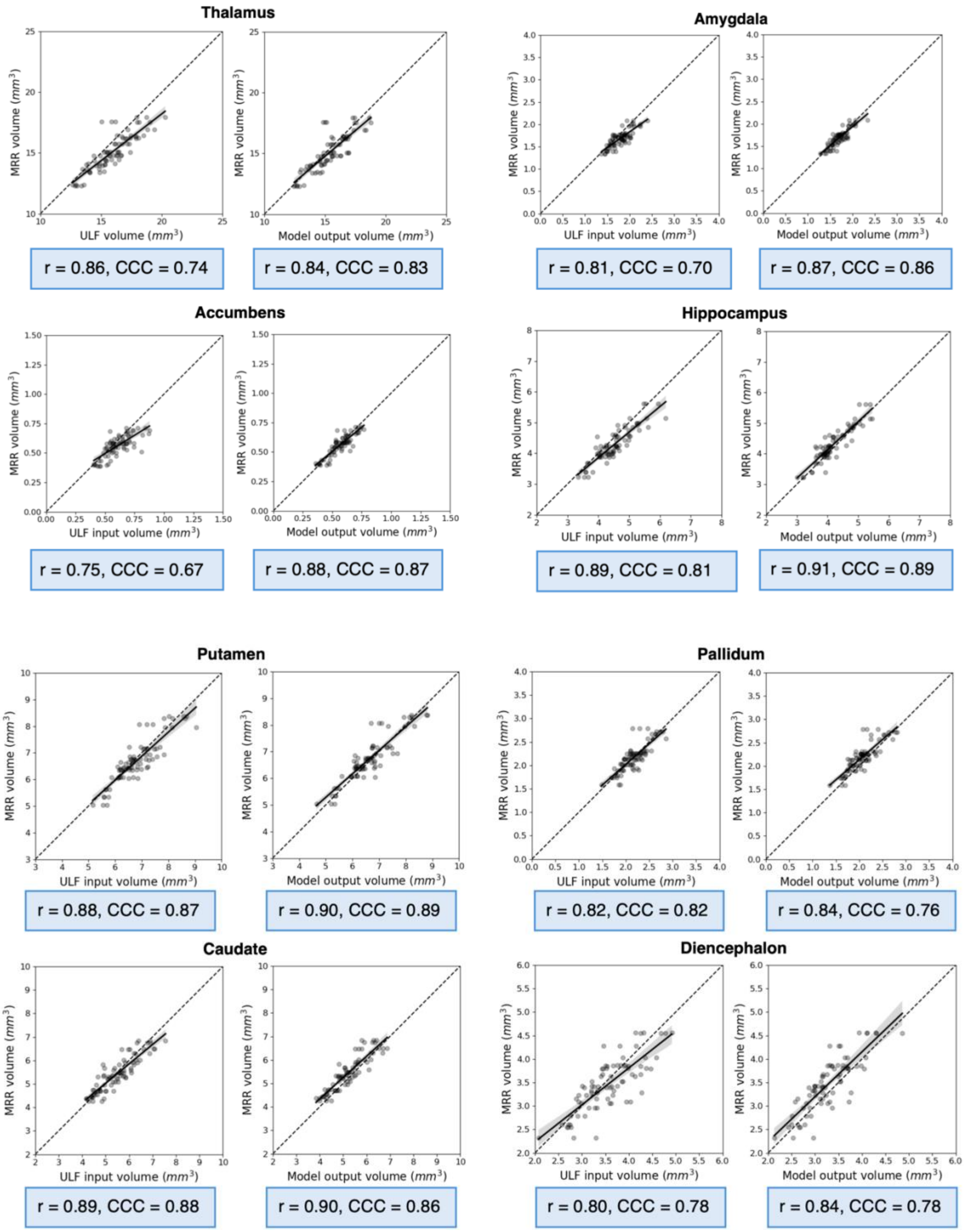


**Figure S4: Tissue volume correlations between MRR and ULF versus model output scans for 8 subcortical regions obtained with BIBSNet.** Blue boxes under the correlation plots report Pearson's correlation coefficient and Lin's concordance correlation coefficient (CCC).

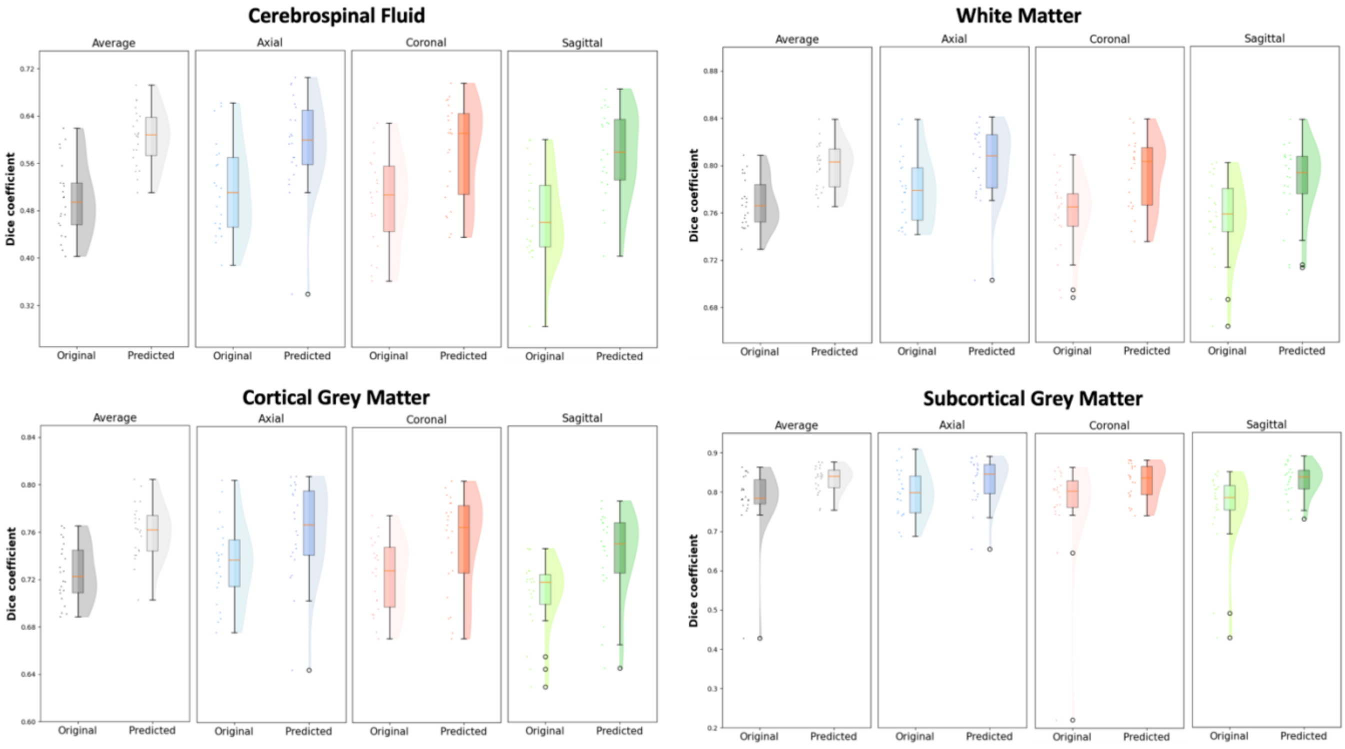


**Figure S5: SynthSeg+ Dice overlap scores between model predictions and MRR scans, and between ULF and MRR scans, for 4 global tissue types.** The ‘All orientations’ plots represent Dice scores obtained by combining axial, coronal and sagittal scans and have 66 data points. ‘Axial’, ‘Coronal’ and ‘Sagittal’ plots represent Dice scores obtained from each orientation separately and have 22 data points each. Original = original ULF scan, Predicted = model prediction.

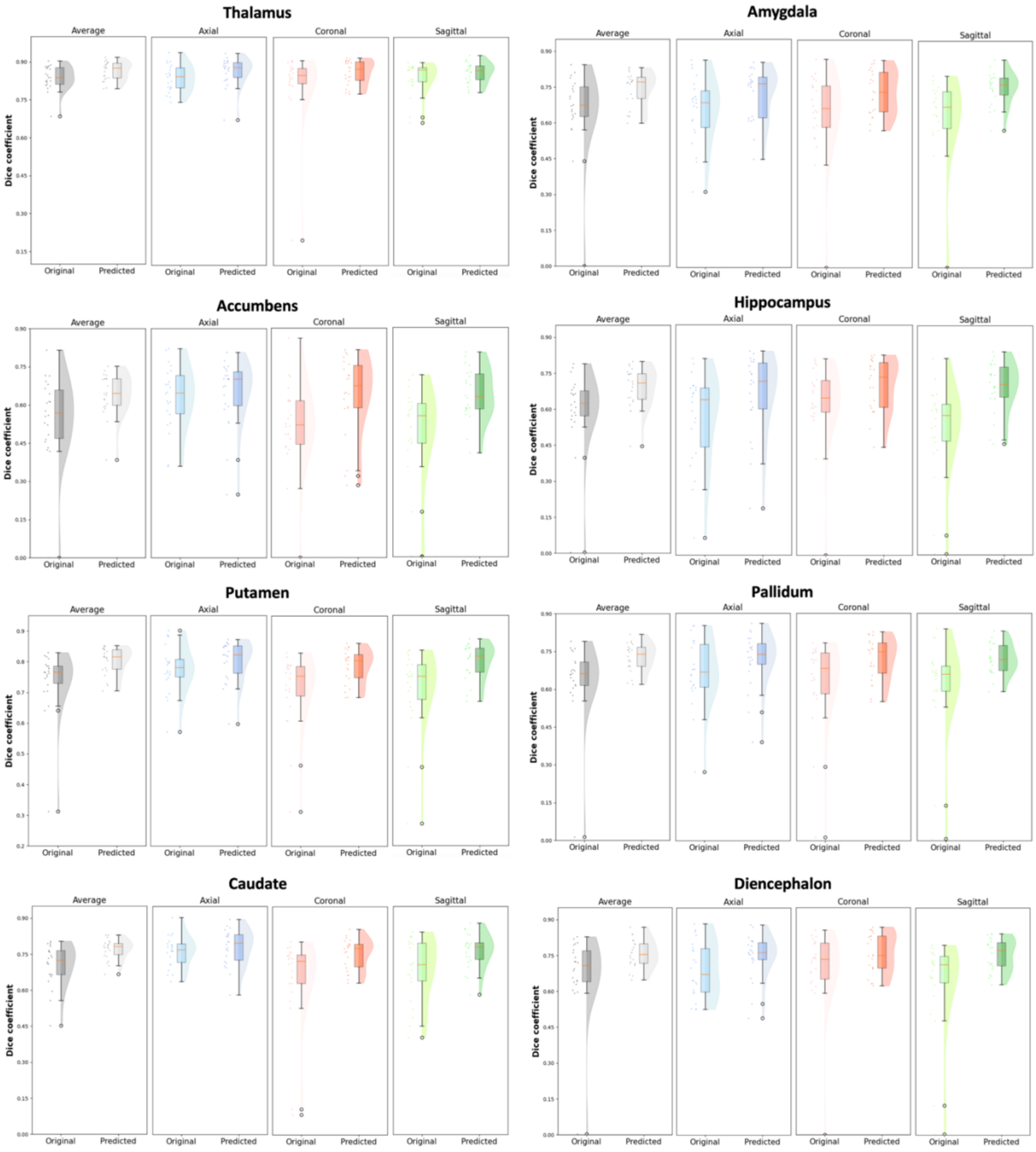


**Figure S6: SynthSeg+ Dice overlap scores between model predictions and MRR scans, and between ULF and MRR scans, for 8 subcortical regions.** The ‘All orientations’ plots represent Dice scores obtained by combining axial, coronal and sagittal scans and have 66 data points. ‘Axial’, ‘Coronal’ and ‘Sagittal’ plots represent Dice scores obtained from each orientation separately and have 22 data points each. Original = original ULF scan, Predicted = model prediction.

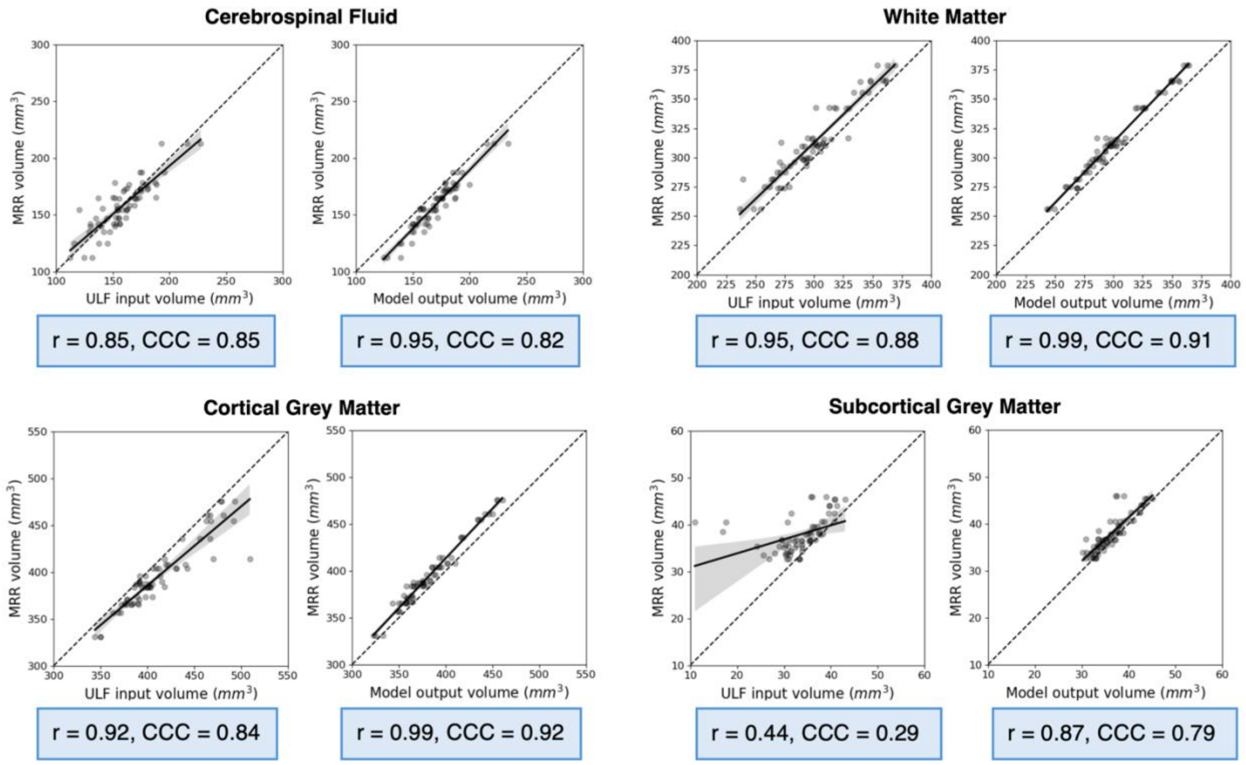


**Figure S7: Tissue volume correlations between MRR and ULF versus model output scans for 4 global tissue types obtained with SynthSeg+.** Blue boxes under the correlation plots report Pearson's correlation coefficient and Lin's concordance correlation coefficient (CCC).

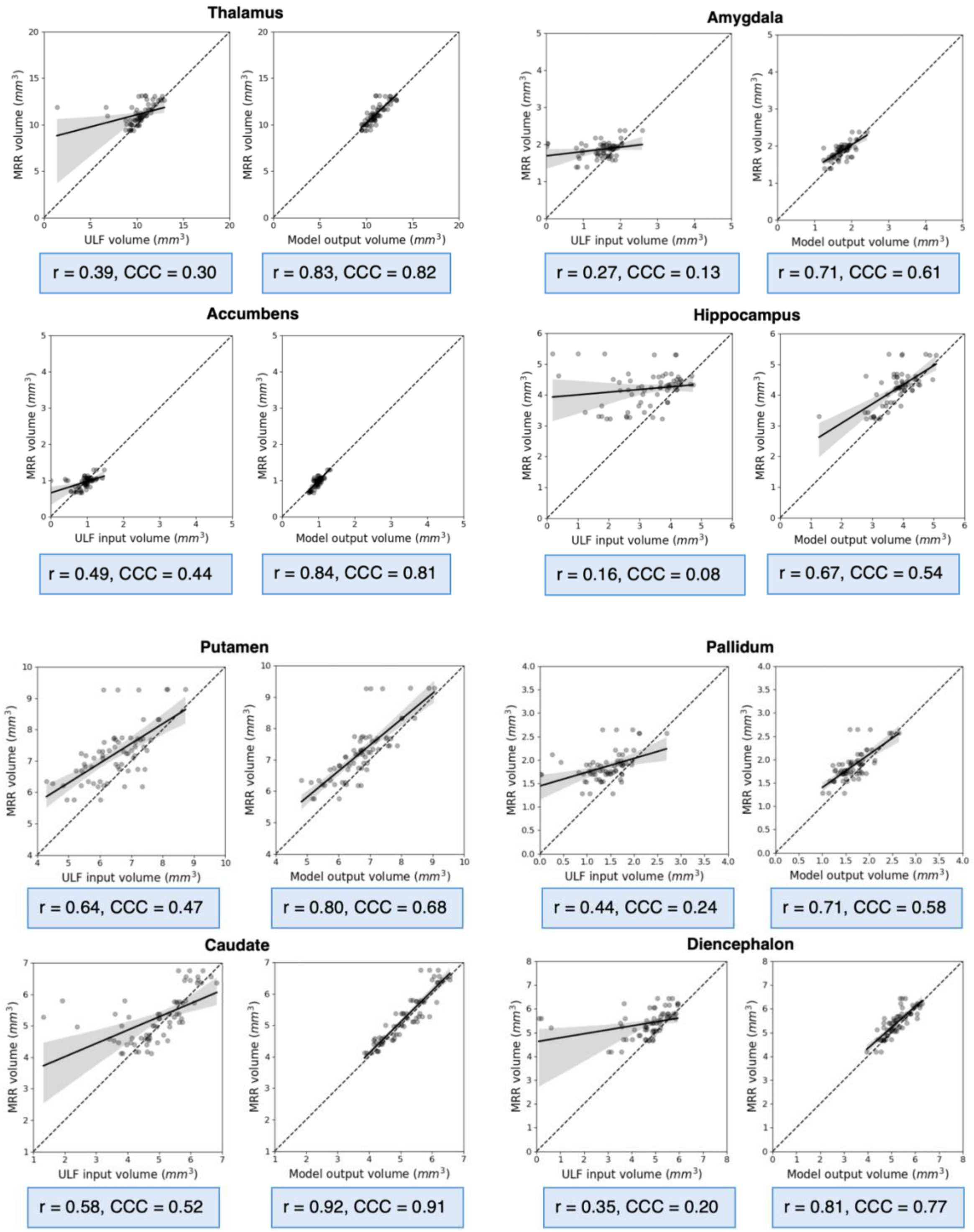


**Figure S8: Tissue volume correlations between MRR and ULF versus model output scans for 8 subcortical regions obtained with SynthSeg+.** Blue boxes under the correlation plots report Pearson's correlation coefficient and Lin's concordance correlation coefficient (CCC).

**Table S1: Image quality metrics for 3 model architectures.** Peak signal-to-noise ratio (PSNR), structural similarity index measure (SSIM), mean absolute error (MAE) and normalized root-mean-squared error (NRMSE) were calculated by comparing model inputs and outputs to reference MRR scans. The model with best performance (in bold) on the validation set was chosen for further optimisation.

| Model Architecture | PSNR (↑) | SSIM (↑) | MAE (↓) | NRMSE (↓) |
|---|---|---|---|---|
| UNet | 22.07 | 0.63 | 0.048 | 0.080 |
| Attention UNet | 18.90 | 0.41 | 0.091 | 0.12 |
| SwinUNETR-v2 | **22.70** | **0.65** | **0.044** | **0.075** |

**Table S2: Image quality metrics for 6 model variants.** Peak signal-to-noise ratio (PSNR), structural similarity index measure (SSIM), mean absolute error (MAE) and normalized root-mean-squared error (NRMSE) were calculated by comparing model inputs and outputs to reference MRR scans. Rank aggregation was first used to determine the optimal loss function (L2 + perceptual), which was subsequently used to compare training on a single- versus multi-orientation input. The model with best performance on the validation set was used for final training.

| | PSNR (↑) | SSIM (↑) | MAE (↓) | NRMSE (↓) |
|---|---|---|---|---|
| | | Loss function | | |
| L1 | 22.70 | 0.65 | **0.044** | 0.075 |
| L2 | **22.90** | 0.63 | 0.045 | **0.073** |
| L1 + perceptual | 22.41 | 0.65 | 0.045 | 0.077 |
| L2 + perceptual | 22.83 | **0.66** | **0.044** | 0.074 |
| | | Input type | | |
| Single-orientation (AXI) | 22.58 | 0.65 | **0.044** | 0.076 |
| Multi-orientation (AXI, COR, SAG) | **22.83** | **0.66** | **0.044** | **0.074** |

**Table S3: Mean SynthSeg+ QC scores.** Mean QC scores for 3 global tissue types and 3 subsets of subcortical regions for each scan type across the 22 participants in the test set. The recommended rejection threshold is 0.65 (Billot et al., 2023), which is on average surpassed for most regions. Since SynthSeg+ was trained on adult scans, we did not reject segmentations based on the automated QC scores but only based on visual QC.

| Region | $AXI_{orig}$ | $COR_{orig}$ | $SAG_{orig}$ | MRR | $AXI_{pred}$ | $COR_{pred}$ | $SAG_{pred}$ |
|---|---|---|---|---|---|---|---|
| CSF | 0.74 | 0.72 | 0.72 | 0.78 | 0.77 | 0.78 | 0.78 |
| WM | 0.75 | 0.73 | 0.74 | 0.78 | 0.78 | 0.77 | 0.78 |
| $GM_{cort}$ | 0.64 | 0.63 | 0.64 | 0.67 | 0.67 | 0.66 | 0.67 |
| Thalamus | 0.80 | 0.77 | 0.77 | 0.83 | 0.83 | 0.83 | 0.83 |
| Putamen + Pallidum | 0.78 | 0.74 | 0.72 | 0.82 | 0.82 | 0.80 | 0.79 |
| Hippocampus + Amygdala | 0.68 | 0.66 | 0.59 | 0.75 | 0.75 | 0.74 | 0.72 |

**Table S4: Wilcoxon signed-rank test applied to Dice scores obtained with SynthSeg+.** Wilcoxon signed-rank test applied to Dice scores between original and predicted scans (all orientations combined and across AXI, COR, and SAG), as well as between predicted Dice scores for each pair of orientations. p-value and RBC are reported for each comparison. RBC is a measure of effect size (strength of the difference between two groups) ranging between -1 and 1, where the closer the RBC is to ±1, the stronger the effect. Abbreviations: RBC = rank biserial correlation, orig. = original, pred. = predicted, CSF = cerebrospinal fluid, WM = white matter, $GM_{cort}$ = cortical grey matter, $GM_{subcort}$ = subcortical grey matter.

| Region | $All_{orig}$ vs $All_{pred}$ (orig. < pred.) | | $AXI_{orig}$ vs $AXI_{pred}$ | | $COR_{orig}$ vs $COR_{pred}$ | | $SAG_{orig}$ vs $SAG_{pred}$ | | $AXI_{pred}$ vs $COR_{pred}$ | | $AXI_{pred}$ vs $SAG_{pred}$ | | $COR_{pred}$ vs $SAG_{pred}$ | |
|---|---|---|---|---|---|---|---|---|---|---|---|---|---|---|
| | p-value | RBC | p-value | RBC | p-value | RBC | p-value | RBC | p-value | RBC | p-value | RBC | p-value | RBC |
| CSF | <.001* | 1.0 | <.001* | 0.75 | <.001* | 0.96 | <.001* | 1.0 | .21 | 0.20 | 0.21 | 0.20 | .35 | 0.10 |
| WM | <.001* | 0.98 | .0037* | 0.64 | <.001* | 0.95 | <.001* | 1.0 | .14 | 0.27 | .078 | 0.35 | .19 | 0.22 |
| $GM_{cort}$ | <.001* | 0.94 | .0026* | 0.66 | <.001* | 0.87 | <.001* | 1.0 | .11 | 0.30 | .078 | 0.35 | .25 | 0.17 |
| $GM_{subcort}$ | .0013* | 0.71 | .015 | 0.53 | <.001* | 0.82 | <.001* | 1.0 | .41 | 0.06 | .38 | 0.08 | .42 | 0.05 |
| Thalamus | .0057* | 0.60 | .013 | 0.54 | .010 | 0.56 | .018 | 0.51 | .25 | 0.17 | .19 | 0.23 | .28 | 0.15 |
| Amygdala | .0018* | 0.68 | .02 | 0.50 | .010 | 0.56 | <.001* | 0.88 | .60 | -0.06 | .85 | -0.25 | .73 | -0.15 |
| Accumbens | .0057* | 0.60 | .25 | 0.17 | .0046* | 0.62 | <.001* | 0.83 | .47 | 0.02 | .27 | 0.15 | .42 | 0.05 |
| Hippocampus | .0013* | 0.71 | .037 | 0.44 | <.001* | 0.83 | <.001* | 0.98 | .87 | -0.28 | .66 | -0.10 | .27 | 0.15 |
| Putamen | <.001* | 0.82 | .064 | 0.38 | <.001* | 0.84 | <.001* | 0.96 | .073 | 0.36 | .29 | 0.14 | .81 | -0.21 |
| Pallidum | <.001* | 0.81 | .037 | 0.44 | <.001* | 0.86 | <.001* | 0.85 | .54 | -0.02 | .34 | 0.11 | .38 | 0.08 |
| Caudate | <.001* | 0.77 | .23 | 0.19 | .0016* | 0.69 | <.001* | 0.81 | .069 | 0.37 | .27 | 0.15 | .55 | -0.03 |
| Diencephalon | .0023* | 0.67 | .023 | 0.49 | <.001* | 0.92 | <.001* | 0.77 | .78 | -0.19 | .49 | 0.01 | .36 | 0.09 |

**Table S5: Volume correlations obtained with SynthSeg+ for 4 global tissue types per orientation.** Volume correlations between MRR and ULF, MRR and model prediction volumes and differences between them (Δ Pearson's r and Δ CCC).

| Region | AXI | | COR | | SAG | |
|---|---|---|---|---|---|---|
| | Pearson's r | CCC | Pearson's r | CCC | Pearson's r | CCC |
| | Volume correlation between MRR and ULF | | | | | |
| CSF | 0.97 | 0.94 | 0.91 | 0.78 | 0.94 | 0.85 |
| WM | 0.97 | 0.94 | 0.93 | 0.81 | 0.97 | 0.89 |
| $GM_{cort}$ | 0.98 | 0.86 | 0.86 | 0.79 | 0.94 | 0.86 |
| $GM_{subcort}$ | 0.63 | 0.49 | 0.37 | 0.24 | 0.48 | 0.25 |
| | Volume correlation between MRR and model prediction | | | | | |
| CSF | 0.98 | 0.91 | 0.98 | 0.88 | 0.95 | 0.70 |
| WM | 0.98 | 0.91 | 0.99 | 0.91 | 0.99 | 0.89 |
| $GM_{cort}$ | 0.99 | 0.93 | 0.99 | 0.93 | 0.99 | 0.90 |
| $GM_{subcort}$ | 0.87 | 0.79 | 0.87 | 0.81 | 0.87 | 0.78 |
| | Δ volume correlation (MRR vs model prediction - MRR vs ULF) | | | | | |
| CSF | 0.01 | -0.03 | 0.07 | 0.10 | 0.01 | -0.15 |
| WM | 0.01 | -0.03 | 0.06 | 0.10 | 0.02 | 0.0 |
| $GM_{cort}$ | 0.01 | 0.07 | 0.13 | 0.14 | 0.05 | 0.04 |
| $GM_{subcort}$ | 0.24 | 0.30 | 0.50 | 0.57 | 0.39 | 0.53 |

**Table S6: Volume correlations obtained with SynthSeg+ for 8 subcortical regions per orientation.** Volume correlations between MRR and ULF, MRR and model predictions and differences between them (Δ Pearson's r and Δ CCC).

| Region | AXI | | COR | | SAG | |
|---|---|---|---|---|---|---|
| | Pearson's r | CCC | Pearson's r | CCC | Pearson's r | CCC |
| | Volume correlation between MRR and ULF | | | | | |
| Thalamus | 0.66 | 0.49 | 0.25 | 0.18 | 0.48 | 0.36 |
| Amygdala | 0.43 | 0.23 | 0.37 | 0.21 | 0.12 | 0.05 |
| Accumbens | 0.78 | 0.77 | 0.48 | 0.43 | 0.43 | 0.34 |
| Hippocampus | 0.26 | 0.17 | 0.09 | 0.06 | 0.17 | 0.05 |
| Putamen | 0.64 | 0.55 | 0.70 | 0.50 | 0.69 | 0.41 |
| Pallidum | 0.64 | 0.48 | 0.45 | 0.25 | 0.45 | 0.17 |
| Caudate | 0.92 | 0.87 | 0.42 | 0.32 | 0.73 | 0.58 |
| Diencephalon | 0.39 | 0.28 | 0.35 | 0.20 | 0.38 | 0.18 |
| | Volume correlation between MRR and model prediction | | | | | |
| Thalamus | 0.84 | 0.82 | 0.82 | 0.81 | 0.83 | 0.82 |
| Amygdala | 0.73 | 0.54 | 0.80 | 0.74 | 0.63 | 0.56 |
| Accumbens | 0.86 | 0.80 | 0.86 | 0.85 | 0.82 | 0.78 |
| Hippocampus | 0.67 | 0.49 | 0.88 | 0.78 | 0.55 | 0.42 |
| Putamen | 0.83 | 0.77 | 0.80 | 0.70 | 0.84 | 0.60 |
| Pallidum | 0.80 | 0.75 | 0.71 | 0.54 | 0.70 | 0.49 |
| Caudate | 0.92 | 0.92 | 0.90 | 0.89 | 0.94 | 0.92 |
| Diencephalon | 0.80 | 0.63 | 0.89 | 0.85 | 0.82 | 0.82 |
| | Δ volume correlation (MRR vs model prediction - MRR vs ULF) | | | | | |
| Thalamus | 0.18 | 0.33 | 0.57 | 0.63 | 0.35 | 0.46 |
| Amygdala | 0.30 | 0.31 | 0.43 | 0.53 | 0.51 | 0.51 |
| Accumbens | 0.08 | 0.03 | 0.38 | 0.51 | 0.39 | 0.44 |
| Hippocampus | 0.31 | 0.32 | 0.79 | 0.72 | 0.38 | 0.37 |
| Putamen | 0.19 | 0.22 | 0.10 | 0.20 | 0.15 | 0.19 |
| Pallidum | 0.16 | 0.27 | 0.26 | 0.29 | 0.25 | 0.32 |
| Caudate | 0.0 | 0.05 | 0.48 | 0.57 | 0.21 | 0.34 |
| Diencephalon | 0.41 | 0.35 | 0.54 | 0.65 | 0.44 | 0.64 |

**Table S7: Image quality metrics for out-of-site external validation set.** Peak signal-to-noise ratio (PSNR), structural similarity index measure (SSIM), mean absolute error (MAE) and normalized root-mean-squared error (NRMSE) were calculated by comparing original ULF and model predictions to reference MRR scans. Best value per metric in bold.

| Comparison | PSNR (↑) | SSIM (↑) | MAE (↓) | NRMSE (↓) |
|---|---|---|---|---|
| ULF vs MRR | 29.41 | **0.90** | **0.013** | 0.035 |
| Model prediction vs MRR | **29.69** | 0.80 | 0.015 | **0.033** |

**Table S8: Median Dice scores on out-of-site external validation set for 4 main tissue types and 8 subcortical regions.** Second column contains values of Dice overlap between ULF and MRR scans and third column contains values of Dice overlap between model predictions and MRR scans. Values correspond to: median obtained with BIBSNet (median obtained with SynthsSeg+).

| Region | ULF | | Model prediction | |
|---|---|---|---|---|
| | BIBSNet | SynthSeg+ | BIBSNet | SynthSeg+ |
| | Main tissue types | | | |
| CSF | 0.83 | 0.45 | 0.83 | 0.53 |
| WM | 0.76 | 0.67 | 0.83 | 0.76 |
| $GM_{cort}$ | 0.76 | 0.67 | 0.75 | 0.73 |
| $GM_{subcort}$ | 0.89 | 0.17 | 0.89 | 0.78 |
| | Subcortical regions | | | |
| Thalamus | 0.90 | 0.17 | 0.89 | 0.87 |
| Amygdala | 0.84 | 0.00 | 0.85 | 0.42 |
| Accumbens | 0.71 | 0.01 | 0.69 | 0.58 |
| Hippocampus | 0.82 | 0.02 | 0.80 | 0.43 |
| Putamen | 0.85 | 0.20 | 0.84 | 0.77 |
| Pallidum | 0.83 | 0.07 | 0.79 | 0.65 |
| Caudate | 0.84 | 0.02 | 0.83 | 0.74 |
| Diencephalon | 0.82 | 0.13 | 0.80 | 0.64 |
